# Zic2 abrogates an alternative Wnt signaling pathway to convert axon attraction into repulsion

**DOI:** 10.1101/759407

**Authors:** Cruz Morenilla-Palao, Maria Teresa López-Cascales, José P. López-Atalaya, Diana Baeza, Luis Calvo-Diaz, Aida Giner de Gracia, Angel Barco, Eloísa Herrera

## Abstract

Wnt signaling is involved in axon pathfinding during brain wiring but it is unknown how Wnt ligands promote attraction or repulsion. In addition, the participation of the canonical (βcatenin-dependent transcription) versus non-canonical (βcatenin-independent) Wnt pathways in this process remains controversial. Here we show that Wnt5a is expressed at the optic chiasm midline and promotes axon crossing by triggering an alternative Wnt pathway that depends on polarized accumulation of βcatenin at the axon terminal but does not activate the canonical pathway. Remarkably, this alternative pathway is silenced by the transcription factor Zic2 in the small subset of ipsilaterally projecting neurons. Zic2 directly regulates genes related to Wnt and Eph signaling that lead to global accumulation of βcatenin but triggers its asymmetric phosphorylation to facilitate the steering of the growth cone. This alternative Wnt pathway found in contralateral axons and its Zic2-mediated abrogation in ipsilateral neurons is likely operating in many other contexts requiring a two-way response to Wnt ligands.

## INTRODUCTION

To establish neuronal connectivity in the adult brain, neurons located far from their targets must extend an axon that navigates over long distances during embryonic development to reach the correct target tissue. Using simple models such as the axonal binary decision of crossing or not the midline in vertebrates or invertebrates, different families of secreted (Netrin, Slits) and membrane (Eph/ephrins) proteins were initially identified as axon guidance molecules (Chédotal, 2019; Herrera et al., 2019; Stoeckli, 2018). Later, the highly conserved proteins Wnts –originally described as key regulators of body axis patterning, cell fate specification, proliferation, cell migration and carcinogenesis– were found to be also involved in axon guidance and mediate both attractive and repulsive responses in different contexts. While Wnt5 repels ventral nerve axons expressing the Derailed receptor in *Drosophila* (Fradkin et al., 1995; Yoshikawa et al., 2003), vertebrate post-commissural axons are attracted by Wnts (Lyuksyutova et al., 2003).

Intracellularly, Wnt signaling is a particularly complex pathway that leads to the activation of two main alternative branches. The canonical and the non-canonical pathway with the latter, in turn, being divided into the Planar Cell Polarity (PCP) and calcium pathways. This classification relies on the participation of βcatenin, an important intracellular signal transducer that links the membrane adhesion protein E-cadherin to the actin cytoskeleton (Jamora and Fuchs, 2002). In the absence of Wnt, βcatenin is constantly degraded by the destruction complex formed by GSK3, Axin and Apc (Apc2 in the nervous system). In the canonical pathway, the binding of Wnt proteins to their receptors (Frizzled and Lrp5/6) triggers the inactivation or disassembly of the destruction complex which reduces βpromotes its accumulation and translocation to the nucleus. There, β forms a complex with Lef/Tcf factors and induces the transcription of specific genes. In contrast, activation of the non-canonical pathway does not depend on βcatenin-driven transcription; instead it relies on changes that affect cytoskeletal organization and calcium homeostasis (Rao and Kühl, 2010). The role of classical PCP proteins, in polarizing epithelial tissues made the non-canonical branch the most likely candidate to translate Wnt signals into the polarization of the motile axon growth cone. Consistent with this view, mutant mice for typical PCP receptors such as Celsr3 (Cadherin EGF LAG seven-pass G-type receptor 3, a.k.a Flamingo) or Vangl2 (Van Gogh-like protein 2), exhibit pathfinding defects in the ascending and descending projections of the brainstem, in the longitudinal axis of the spinal cord and in other fiber tracts (Chai et al., 2014; Fenstermaker et al., 2010; Shafer et al., 2011; Tissir et al., 2005). However, more recent studies have shown that manipulation of different components of the canonical pathway, including βcatenin, alter axon midline behavior in the chick spinal cord (Avilés and Stoeckli, 2016). This, along with studies showing that Wnt5a may act as an activator of the canonical pathway (Mikels and Nusse, 2006), has challenged the idea of PCP as the only Wnt signaling branch involved in guidance. Therefore, the mechanisms by which Wnt proteins mediate axon attraction or repulsion as well as the branch of the Wnt pathway involved in axon guidance remain unclear.

In the developing visual system, retinal ganglion cells (RGCs) projecting ipsilaterally (iRGCs) are located in the peripheral ventro-temporal retina and contralaterally projecting ganglion cells (cRGCs) distribute in the remainder retina. The spatial segregation of iRGCs and cRGCs facilitates their genetic manipulation and the labeling and visualization of individual axons when they are crossing or avoiding the midline. By using this binary system, here we disentangle some of the unknown mechanisms underlying Wnt signaling pathways during axon guidance decisions. We first demonstrate that cRGC and iRGC axons transduce Wnt5a signaling differentially. In contralateral neurons, a novel form of Wnt signaling that depends on accumulation of βcatenin but is non-canonical, is activated to promote midline crossing. In ipsilateral axons, the positive response to Wnt5a is switched off by Zic2, a transcription factor (TF) previously described as the determinant of ipsilateral identity and known to induce the expression of the tyrosine kinase receptor EphB1 (Escalante et al., 2013; García-Frigola et al., 2008; Herrera et al., 2003). Then, we identified the set of genes directly regulated by Zic2 that silence the positive response to Wnt5a. Concomitant with the Zic2-induced blockage of Wnt5a positive signaling, ipsilateral axons undergo EphB1-βcatenin at the growth cone to trigger asymmetric destabilization of actin filaments and facilitate axon steering.

## RESULTS

### Wnt5a enhances the growth of contralaterally projecting axons

Wnt signaling is known to play an important role in axonal navigation in different contexts and species, but its function in the guidance of visual axons at the midline has not been explored. To assess the role of Wnt signaling in the navigation of retinal axons, we first analyzed the expression of different members of the Wnt family at the optic chiasm by *in situ* hybridization. Among the different Wnts expressed at the chiasm region (**Figure S1**), we found that *Wnt5a* is expressed at the midline in a spatiotemporal pattern that resembles that of *ephrinB2* (**Figure 1A**), a repulsive guidance molecule expressed by glial cells that induces the turning of iRGCs (Williams et al., 2003). This result points towards Wnt5a being a candidate molecule responsible for mediating axon guidance at the midline.

**Figure 1.**
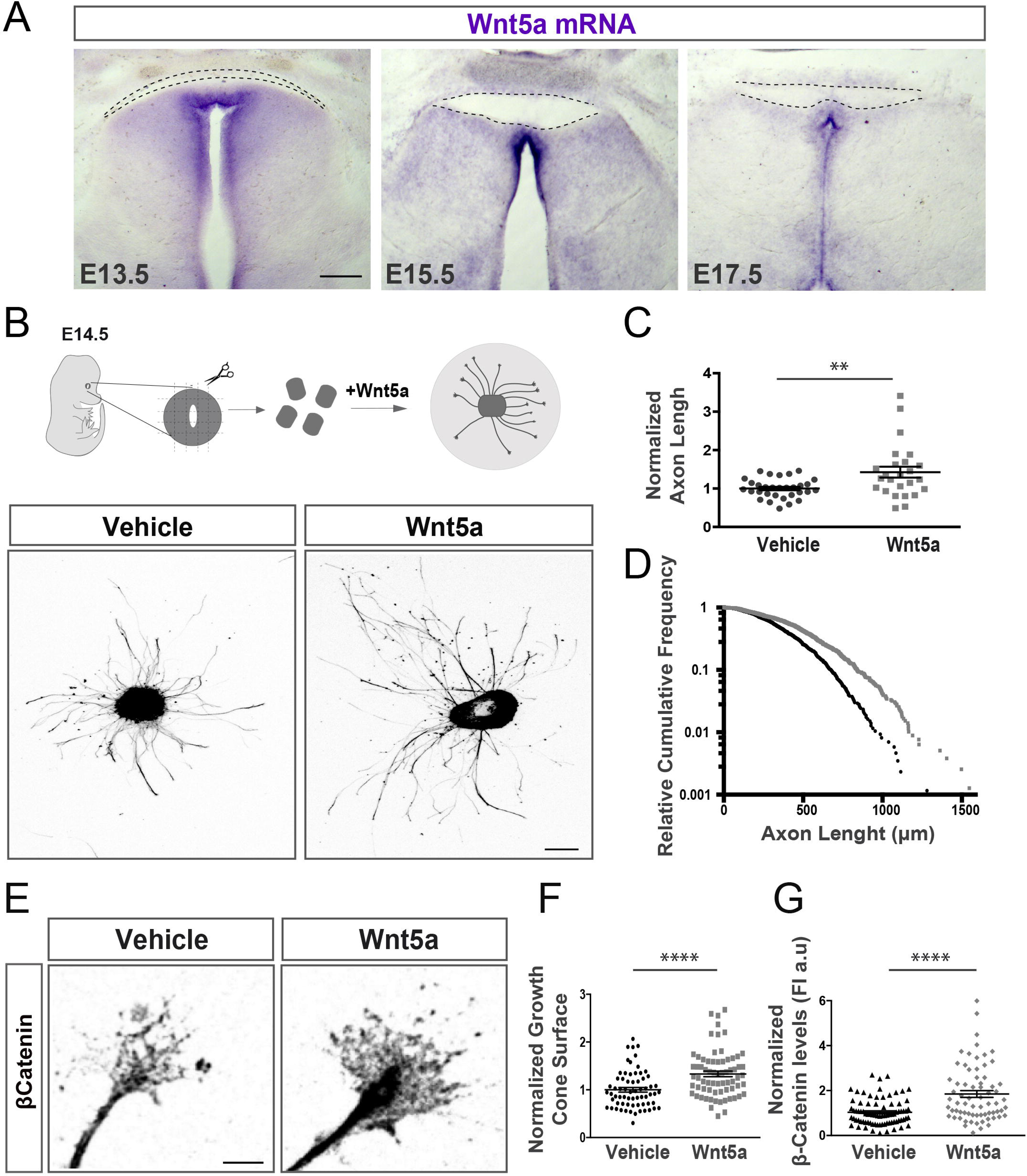
Wnt5A is expressed at the midline and enhances axonal growth of contralateral axons. **(A)** In situ hybridization for Wnt5a in coronal sections from E13.5, E15.5 and E17.5 embryos. Scale bar: 20μM. **(B)** β-tubulin staining on retinal explants from E14.5 embryos cultured for 24 hours. Retinal explants from the central retina treated with recombinant Wnt5a protein display longer neurites compared with vehicle-treated explants. Scale bar: 100 μM. **(C)** Quantification of axonal length in explants treated with vehicle versus Wnt5a. Axonal length values in Wnt5a-treated explants (n=25) were normalized to the mean value of axons treated with the vehicle (n=31). n=number of explants (Two-tailed Mann-Witney U test, p=0.007). **(D)** Relative cumulative frequency histogram showing axonal length in explants growing in the presence of Wnt5a (gray, n=927) or the vehicle (black, n=861). Each dot represents an individual neurite. n=number of axons from three biological replicates. **(E)** Immunohistochemistry for βcatenin in the growth cone of ganglion cells form contralateral axons treated for 1 hour with Wnt5a or vehicle. Note that Wnt5a-treated cones occupy a larger area and show higher levels of βcatenin. Scale bar: 5μM. **(F, G)** Quantification of the growth cone surface (vehicle n=66; Wnt5a n=69) and the levels of βcatenin (vehicle n=81; Wnt5a n=69) in the cone of Wnt5a and vehicle treated retinal explants. Individual axon length and values for βcatenin fluorescence intensity (FI) from Wnt5a-treated growth cones were normalized to vehicle mean. βcatenin levels Two-tailed unpaired t-test, P<0.0001. Results show mean ± SEM.

To investigate whether Wnt5a has an effect on RGC axons, explants from the central retina of E14.5 mouse embryos (i.e. embryos containing contralaterally but not ipsilaterally projecting neurons) were cultured for 12 hours in the presence of Wnt5a. The axons from explants co-cultured with Wnt5a extended significantly longer axons than those not exposed to Wnt5a (**Figure 1B–D**). These results indicate that RGC axons respond positively to Wnt5a, confirming the participation of Wnt signaling in visual axons pathfinding.

### A non-canonical but βcatenin-dependent branch of the Wnt pathway mediates midline crossing

Intriguingly, the experiments with retinal explants described above revealed a significant increase in both the levels of βcatenin and the size of the growth cone after acute exposure of the explant to Wnt5a (**Figure 1E–G**). To directly assess the participation of βcatenin in midline crossing, we downregulated βcatenin in RGCs at the time that the majority of visual axons are crossing the midline. Short hairpin RNA against βcatenin (*Ctnnb shRNA)* was injected and electroporated into the retinas of E13.5 mouse embryos and the axonal projection phenotype of targeted neurons at the optic chiasm was analyzed five days later. As a control, retinas were also electroporated with plasmids bearing random *shRNAs*. In addition, plasmids encoding EGFP were co-injected in both cases to visualize the axons of targeted neurons at the chiasm. While the axons of neurons expressing random *shRNAs* crossed the midline, those electroporated with *Ctnnb shRNA* plasmids showed strongly altered axonal trajectories at the midline. While a subpopulation of these axons took the ipsilateral path, most of them stalled at the midline (**Figure 2A–B**). These results are consistent with previous findings showing midline defects of commissural spinal axons after downregulation of βcatenin (Avilés and Stoeckli, 2016).

**Figure 2.**
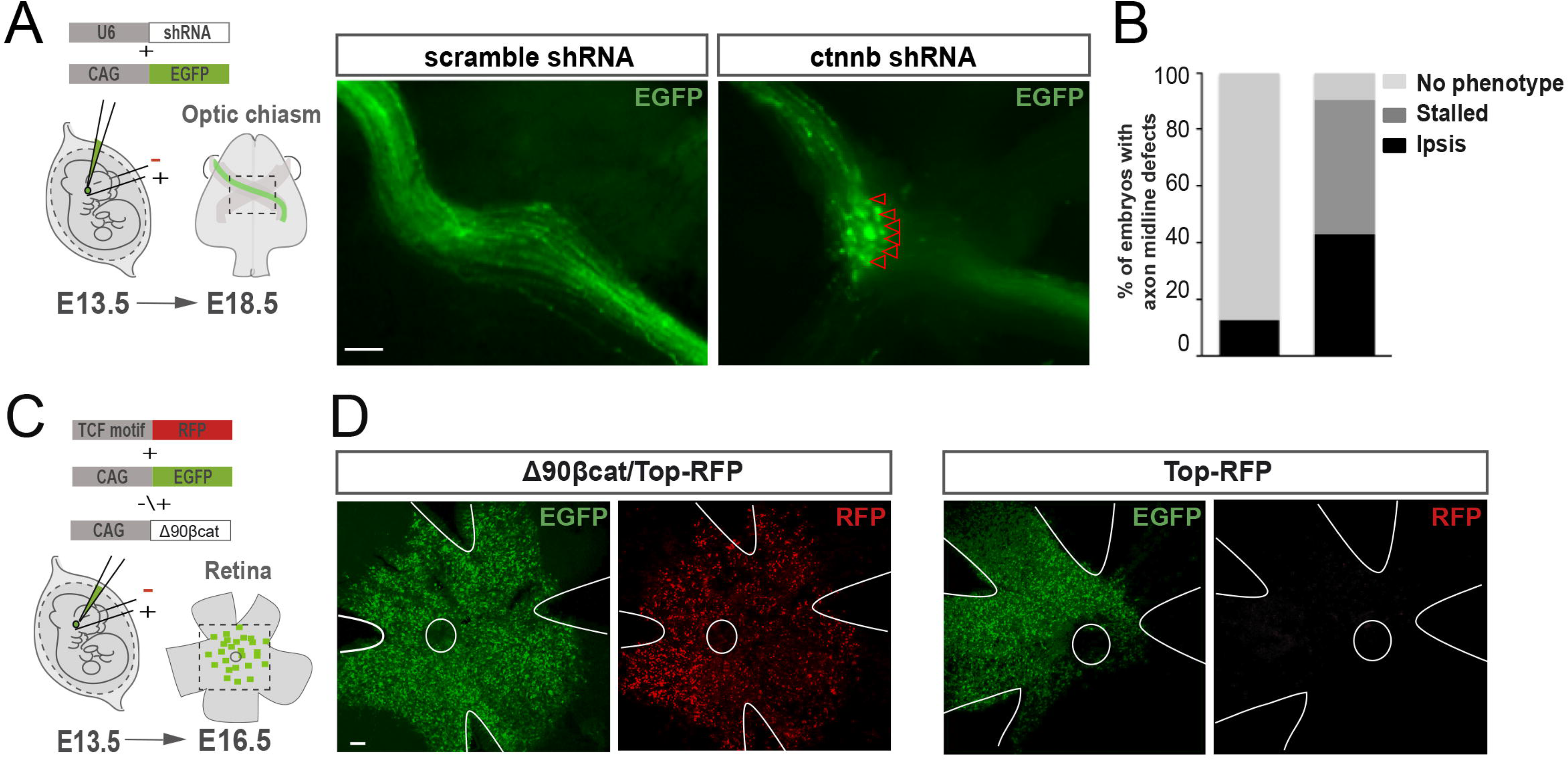
βcatenin is necessary for midline crossing. **(A)** Plasmids encoding EGFP plus control shRNA or shRNA against βcatenin (*Ctnnb shRNAs)* were electroporated into one eye of E13.5 embryos and EGFP-axons into the optic chiasm were analyzed at E18.5. Right panels are representative images of optic chiasms from embryos electroporated with Ctnnb shRNA or scramble shRNAs plasmids. Downregulation of βcatenin in contralaterally projecting neurons inhibits midline crossing. **(B)** Graphs represent the percent of embryos showing midline crossing defects after electroporation with scramble shRNA (n=10; No Phenotype (NP)=87.5%, Ectopic Ipsilateral projection (Ipsi)=12,5%) or Ctnnb shRNAs (n=42; NP=9.52%; Ipsi= 42.86% and Stalled=47.62%) plasmids. **(C)** Summary of the experimental approach. **(D)** Representative immunohistochemistry for RFP and EGFP in wholemount E16.5 retinas electroporated at E13.5 with Top-RF- and EGFP-, or Δ90-βCatenin /Top-RFP- and EGFP-encoding plasmids show that canonical Wnt signaling is not activated in cRGCs at the time that visual axons transverse the midline. The experiment was repeated at least 3 times for each condition with similar results.

The effects of Wnt5a treatment and the phenotype obtained after knocking down βcatenin on cRGCs was suggesting that contralateral neurons could be activating the canonical pathway while confronting Wnt5a at the chiasm. To test this hypothesis, we electroporated retinas from E13.5 embryos with a reporter plasmid for canonical Wnt signaling (referred to as Top-RFP) that contains several TCF/LEF binding sites upstream of the coding sequence of the red fluorescence protein (Rabadán et al., 2016). As a positive control, we used an N-terminal truncated form of βcatenin (Δ90βcat) that lacks of phosphorylation sites responsible for its degradation and tends to accumulate intracellularly promoting βcatenin-dependent transcription (Wrobel et al., 2007). Retinas from embryos electroporated with Top-RFP plus Δ90βcat expressed high levels of RFP two days later. However, no red signal was detected in retinas electroporated with Top-RFP alone (**Figure 2C, D**), indicating that although βcatenin is required for midline crossing, canonical Wnt signaling is not active in RGCs when the axons cross the chiasm.

### Zic2 silences the positive response to Wnt5a

Above we demonstrated that Wnt5a enhances the growth of contralateral axons and triggers the local accumulation of βcatenin at the tip of the cone. Would ipsilateral axons exhibit a different behavior? Given that the number of iRGCs concentrated at the ventrotemporal retina is too small to prepare explants free of cRGCs, we tested our hypothesis by comparing the axonal response of RGCs electroporated or not with Zic2, the TF that specifies ipsilaterally projecting neurons (Escalante et al., 2013; Herrera et al., 2003). In this model, Zic2-positive RGCs functionally correspond to iRGCs. Retinal explants from embryos electroporated with EGFP alone or with Zic2 plus EGFP were plated on dishes and incubated with Wnt5a for 12 hours (**Figure 3A**). As expected, the axons of EGFP-electroporated explants (controls) extended significantly longer axons in the presence of Wnt5a than vehicle (**Figure 3B, similar to Figure 1C**). In contrast, the axons from Zic2-expressing explants did not exhibit enhanced growth after Wnt5a treatment (**Figure 3A–B**), indicating that Zic2 is able to modify the response of contralateral axons to Wnt5a. Moreover, the acute exposure of Zic2-expressing neurons to Wnt5a resulted in a significant reduction in the size of their growth cones compared to controls (**Figure 3C–D**), indicating that, although they do not respond by enhancing their growth, Zic2 RGC axons do sense Wnt5a.

**Figure 3.**
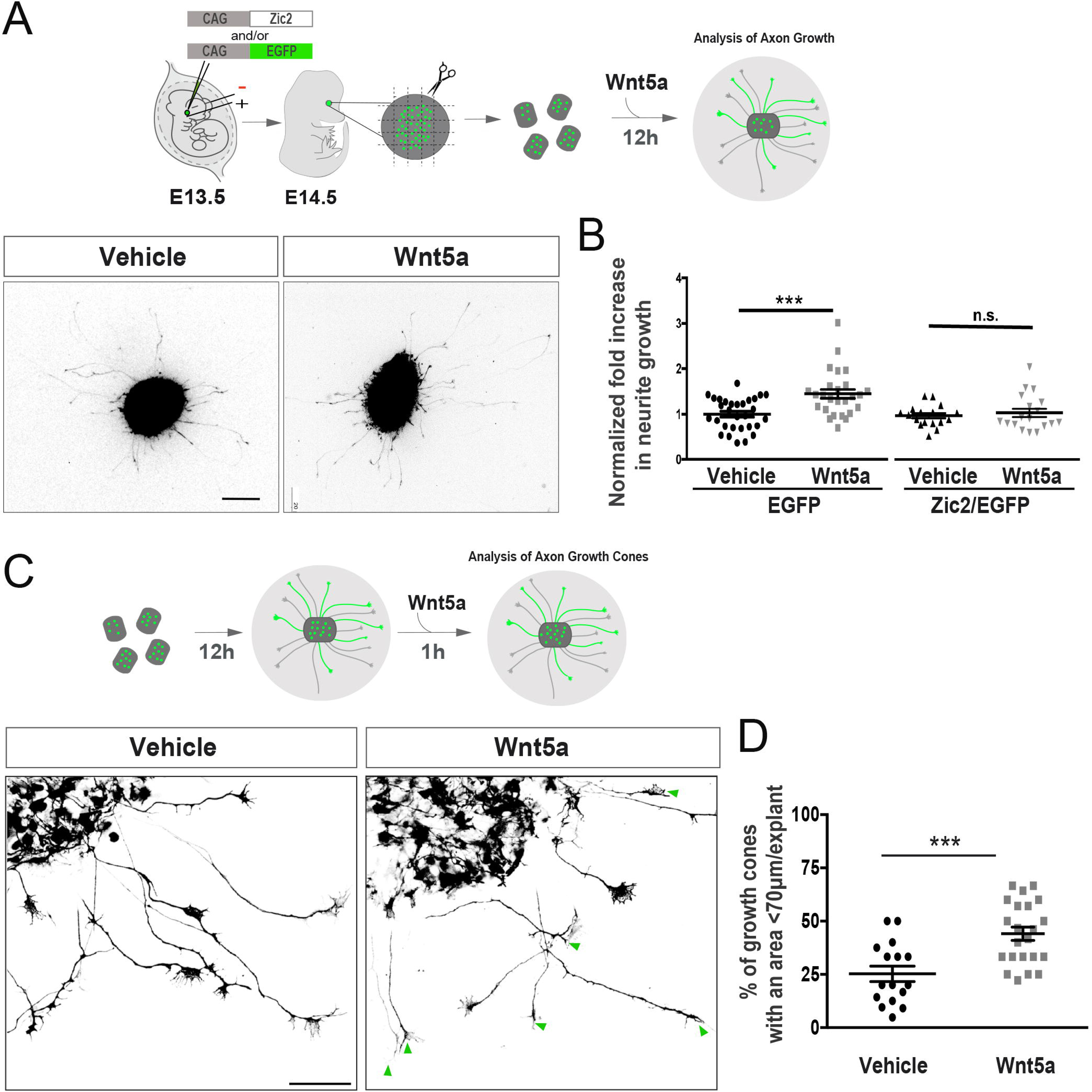
The axons of Zic2-expressing neurons do not show a positive response to Wnt5a. **(A)** Upper panels, show the experimental approach used to assess the response of Zic2 expressing-neurons to Wnt5a. Retinal explants from E14.5 embryos electroporated at E13.5 with EGFP- or Zic2/EGFP-encoding plasmids were isolated and cultured with or without Wnt5a. The bottom images are retinal explants from electroporated embryos incubated with Wnt5a or vehicle. Scale bar: 200μM **(B)** Quantification of axonal length in axons from RGCs expressing EGFP (Vehicle n=30, Wnt5a n=26) or Zic2/EGFP (Vehicle n=17, Wnt5a n=18) grown in the presence of Wnt5a or vehicle. In Wnt5a-treated explants, axon length values were normalized to the mean value of the axons in explants treated with the vehicle. Two-tailed unpaired t-test, EGFP p-value=0.003, Zic2 p-value=0.563. **(C)** Schema summarizing the experimental approach. Images show axons from representative retinal explants isolated from electroporated embryos that were cultured for 24 hours and exposed to Wnt5a or vehicle for 1 hour. Green arrow heads point out axons with growth cones smaller than 70 microns. Scale bar: 50μM **(D)** Quantification showing the percentage of growth cones smaller than 70 microns in Zic2-expressing explants incubated with Wnt5a or the vehicle (Vehicle n=16; Wnt5a n=22; P=0.0004) (Two-tailed unpaired t-test *** p<0.001). Results from three biological replicates. Results show mean ± SEM.

### Zic2 regulates many genes related to the Wnt signaling in RGC axons

Our results show that cRGCs and Zic2-iRGCs respond differentially to Wnt5a. To unveil the mechanisms underlying this change in the axonal response, we analyzed the transcriptome of Zic2-transduced RGCs. Retinas from E13.5 embryos electroporated with Zic2 plus EGFP-encoding plasmids or with EGFP plasmids alone, were isolated and EGFP cells sorted by flow cytometry 36 hours after electroporation. Then, the transcriptomes of EGFP or Zic2/EGFP cells were compared (**Figure 4A and S2A**). This mRNA-seq screen identified 423 upregulated genes and 192 downregulated genes (P_adj_ < 0.1) (**Figure 4B**). As expected, Zic2 was the most upregulated gene (**Figure S2B**). Genes encoding for the tyrosine receptor EphB1 and the serotonin transporter Sert – previously described as Zic2 targets (García-Frigola et al., 2008; García-Frigola and Herrera, 2010) – were also upregulated (**Figure S2C**). In addition, Zic2 expression strongly downregulated Sox4, a TF that promotes cRGC differentiation and axonal midline crossing (Kuwajima et al., 2017) (**Figure 4B and Figure 5H**), as well as Brn3a and Isl2 which have been reported to be expressed in contralateral but not in ipsilaterally projecting neurons (Pak et al., 2004; Quina, 2005) (**Figure S2D**). Consistent with these results, Gene Ontology (GO) enrichment analysis for differentially expressed genes revealed that upregulated genes were associated with biological processes such as *Neuron projection* and *Cell-cell signaling*, which are essential processes in axon guidance. Furthermore, the most enriched *Cellular Component* term was *Axon* (**Figure 4C**).

**Figure 4.**
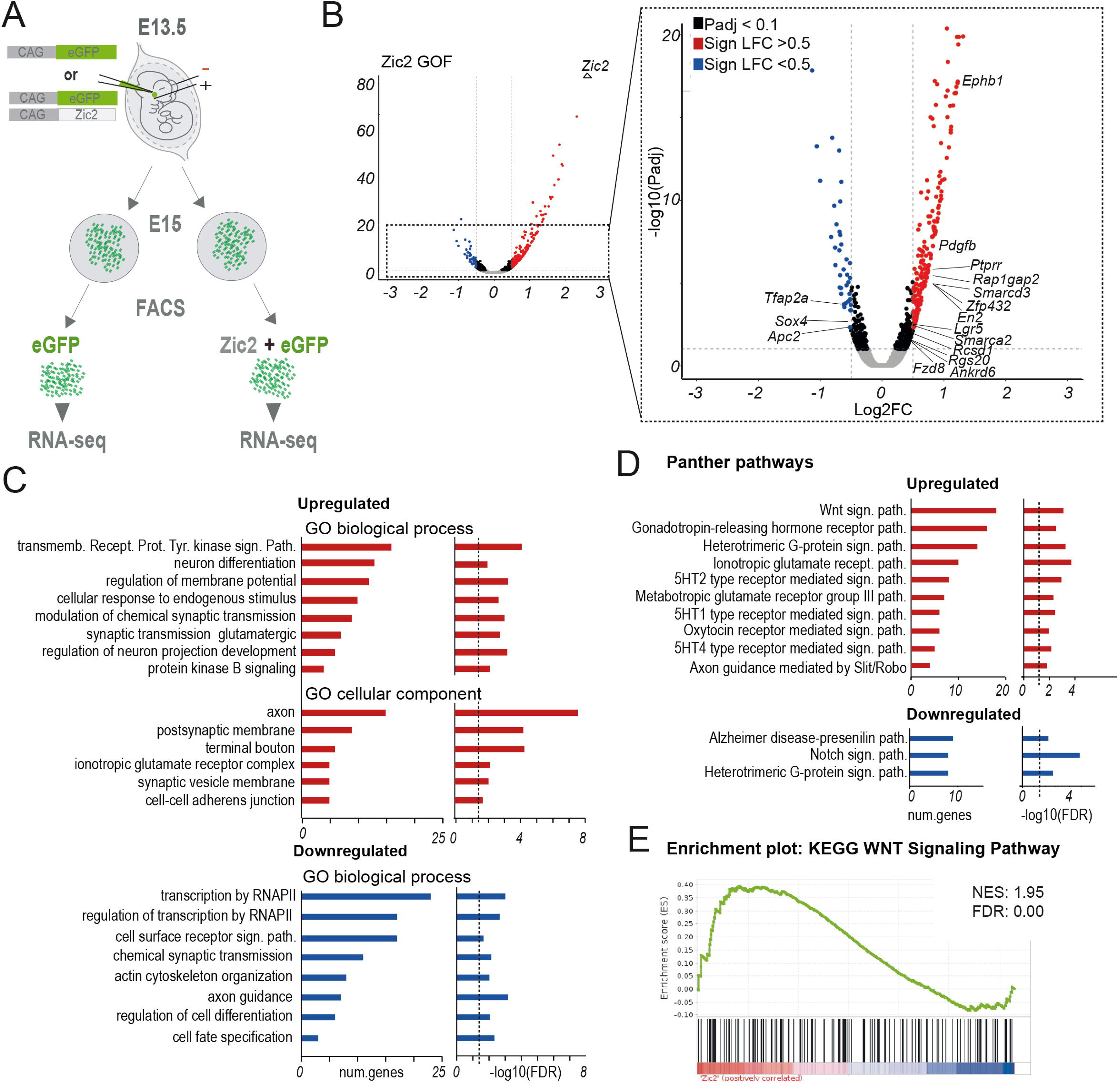
Transcriptome analysis in RGCs transduced with Zic2. **(A)** Scheme representing the experimental design of the RNA-seq screen. **(B)** Volcano plot showing differentially expressed genes (DEGs) 36 hours after Zic2 electroporation in contralateral RGCs. The name of relevant DEGs is indicated in the amplification inset. The significance value for the change in Zic2 is above the scale (P-adj = 0). **(C)** GO enrichment analysis of genes upregulated (top graphs) and downregulated (bottom graphs) after Zic2-induced expression in RGCs. The bar graphs present the significance of the enrichment (right) and the number of genes involved (left). **(D)** Panther pathway enrichment analysis of genes upregulated (top graphs) and downregulated (bottom graphs) after Zic2-induced expression in RGCs. The bar graphs present the significance of the enrichment (right) and the number of genes involved (left). **(E)** GSEA of DEGs after Zic2-transduction of RGCs detects a non-random distribution of genes involved in Wnt signaling. The normalized enrichment score (NES) and FDR values are shown in the upper right corner of the graph.

**Figure 5.**
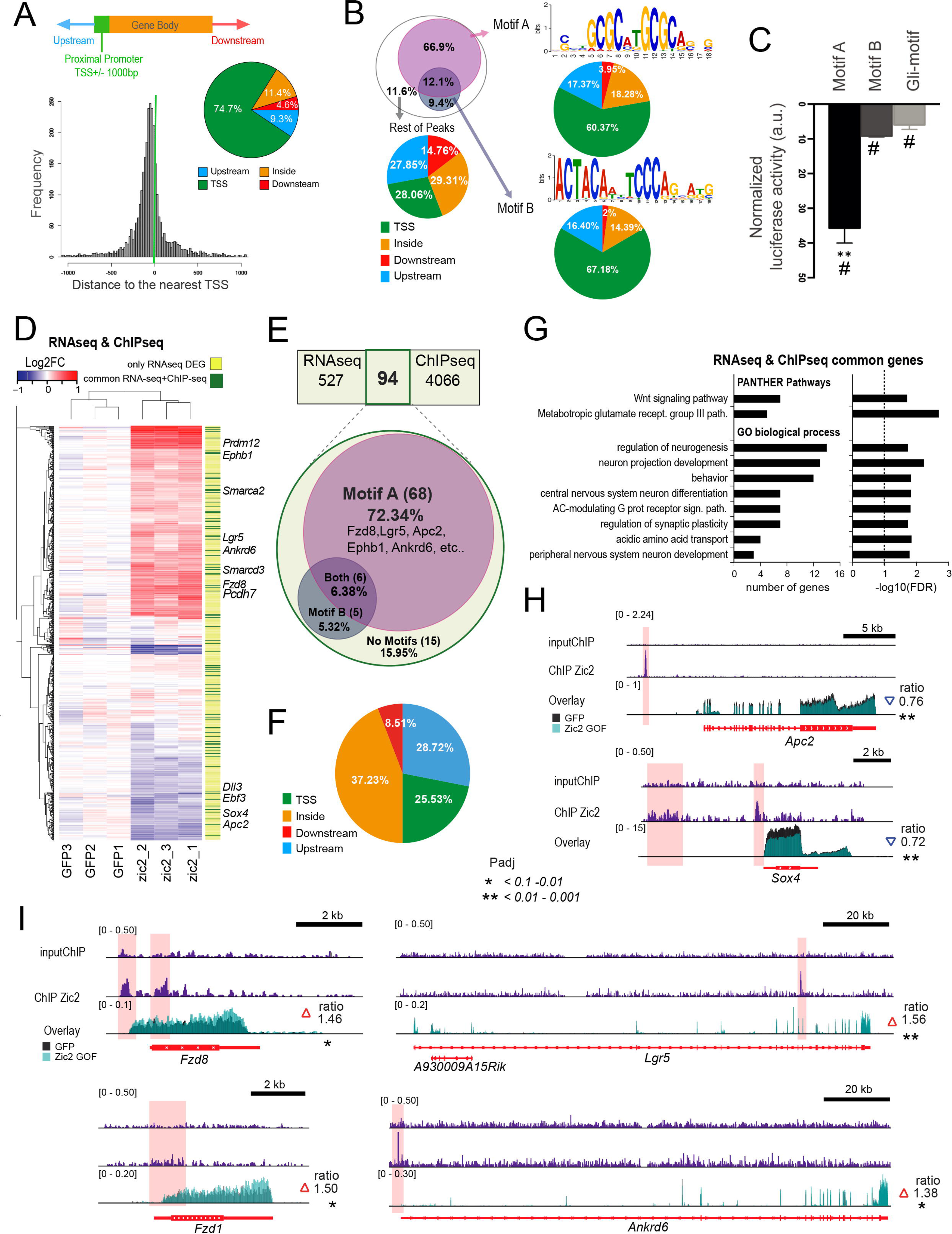
Zic2 regulates Wnt signaling genes. **(A)** Distribution of Zic2 peaks across gene features. Most peaks locate in the proximity of the transcription start site (TSS). **(B)** Unbiased identification of TF binding motifs enriched in Zic2 peaks. The Venn diagram shows the co-presence of the two main motifs detected in our screen. The three sector graphs show the distribution across gene features of the peaks containing motif A, motif B and lacking both motifs. **(C)** Luciferase assay in HEK cells showing a stronger transactivation of the reporter gene driven by Zic2 recruitment by motif A than by motif B or Gli. Note that the three motifs produce a significant increase when compared to the control situation which is the reporter plasmid lacking the motifs (two-tailed unpaired t-test; *p<0.05; **p<0.01). Data from three independent experiments were analyzed. Results show mean ± SEM. **(D)** Heatmap of DEGs detected in our RNA-seq screen. The right column (yellow) highlights the genes detected in both the RNA-seq differential expression screen and the Chip-seq screen for Zic2-bound genes (in black). **(E)** Distribution of motif A and B in the common gene set. **(F)** Distribution of Zic2 binding across gene features in the common gene set. **(G)** GO and PANTHER enrichment analyses of the common gene set. The bar graphs present the significance of the enrichment (right) and the number of genes involved (left) **(H)** ChIP-seq and RNA-seq profiles for two relevant Wnt signaling genes downregulated by Zic2. **(I)** ChIP-seq and RNA-seq profiles for four relevant Wnt signaling genes upregulated by Zic2.

To identify the main signaling pathways affected by Zic2 expression, we conducted a PANTHER pathways analysis that revealed a significant enrichment for genes involved in the Wnt signaling pathway among the upregulated genes (**Figure 4D**). Gene set enrichment analysis (GSEA) confirmed the significant impact of Zic2 expression on Wnt pathway genes and revealed that Wnt-related genes are also found in the set of genes downregulated by Zic2 (**Figure 4E**).

To define the gene set directly regulated by Zic2 in axon guidance processes, we performed chromatin immunoprecipitation assays followed by massive sequencing (ChIP-seq) using a specific Zic2 antibody. Since the population of Zic2-expressing RGCs is extremely low (we had to use more than 170 embryos per sample), we additionally prepared chromatin from dorsal spinal cord neurons (dSCNs), which also project ipsilaterally upon Zic2 expression (Escalante et al., 2013) but are much more abundant than iRGCs. The comparison of both profiles showed that although Zic2 peaks are more prominent in dSCNs, likely reflecting the higher number of Zic2 cells in the spinal cord, the occupancy profiles in iRGCs and dSCN are very similar (**Figure S3A**). This is true with the exception of a few genes, such as EphB1 and EphA4, which are specific for iRGCs and dSCN, respectively (**Figure S3B**). These latter results are consistent with our previous studies demonstrating that Zic2 controls the expression of EphB1 in RGCs and EphA4 in dSCN (Escalante et al., 2013; García-Frigola et al., 2008).

The mapping of Zic2 peaks revealed that Zic2 primarily binds to promoter regions. Although previous analyses in embryonic stem cells reported that Zic2 binds preferentially to enhancer regions (Luo et al., 2015), our results in the chromatin of differentiated neurons indicate that almost 75% of the peaks map in the proximity of transcription start sites (TSS) (<1 kb upstream) (**Figure 5A**) and only 13.9% of the peaks locate into putative enhancers (**Figure 5B**). These results denote fundamental differences either in the binding profile of Zic2 in stem cells and postmitotic neurons.

Intriguingly, the search for motif in Zic2-bound regions using the MEME suite retrieved two consensus binding motifs, which we refer to as motif A and motif B (**Figure 5B**). Motif A resembles the consensus motif recognized by NRF1 (JASPAR: MA0506.1) and has not been previously associated with Zic2 or any other member of the Zic family, while motif B is similar to a sequence described as a secondary Zic2 binding motif (Ishiguro et al., 2018; Sakai-Kato et al., 2008). Motif A was found in 79% of the Zic2 peaks, whereas motif B was only found in 21.5% of the peaks. Zic2 binding to both motifs was detected in 12.1% of the loci (**Figure 5B**). Luciferase assays in HEK293 cells transfected with Zic2 in conjunction with plasmids bearing the thymidine-kinase basal promoter-β-Gal driven by motif A or motif B (see the Material and Methods section), revealed that while Zic2 may interact with both motifs, it shows much higher affinity for motif A. As a control, since Zic TFs have been proposed to antagonize Gli TFs and compete for the same binding sites (Mizugishi et al., 2001), we also analyzed Zic2 binding to the Gli binding motif and found a similar affinity as for motif B (**Figure 5C**). Altogether, these results indicate that motif A enables the strongest transactivation by Zic2 recruitment.

Consistent with our mRNA-seq data, GO analysis of Zic2-bound genes in differentiated neurons revealed a significant enrichment for the Wnt signaling pathway (**Figure S3C**). Moreover, Wnt signaling was also the most enriched pathway among the genes that were both detected as Zic2-bound in the ChIP-seq screen and differentially expressed in the mRNA-seq screen (**Figure 5D-G**). The overlapping gene set (94 genes; Table S1) includes a 37% of downregulated genes and 63% of upregulated genes and Wnt-related Zic2 targets are found in both groups. Key components of the pathway, such as *Apc2* or *Sox4*, are downregulated (**Figure 5H**), while others, such as *Fzd1, Fzd8, Lgr5, En2 or Diversin/Ankrd6*, are upregulated (**Figure 5I**) in Zic2-expressing retinal cells, suggesting that Zic2 may act as an activator or a repressor depending on the genomic context, likely forming complexes with other TFs. Notably, Zic2 peaks were found at the promoter (25%) or intronic putative enhancers (37%) of these *bona fide* target genes, and 80% of them included motif A in the regulatory sequences (**Figure 5E, F and Figure S3**). Remarkably, NRF1, the TF that also recognizes the motif A found in Zic2 peaks, has been proposed to interfere with the Wnt canonical pathway in *Drosophila* (Xin et al., 2011). Interestingly, other Wnt-related genes such as *Ctnnb1* (βcatenin), *Ryk*, *Rhoa*, *Gsk3b*, *Pias4*, *Rock1, Dvl1, Dv2* and *Axin1*, which showed strong Zic2 binding to their promoters, did not change their expression upon Zic2 induction (**Figure S3**).

Overall, these genome-wide analyses show that Zic2 regulates very similar genetic programs in different types of ipsilaterally projecting neurons and underscore a strong link between Zic2 and the Wnt pathway.

### The gene network triggered by Zic2 promotes accumulation of βcatenin

To further define the molecular mechanism underlying the control of Zic2 in the Wnt pathway, we sought to validate our genomic screens by performing *in situ* hybridization (ISH) and immunostaining in retinal sections for some of the candidate genes retrieved in both screens (RNA-seq + ChIP-seq). Thus, ISH for *Fzd8 mRNA* demonstrated a restricted expression in the ventrotemporal retina coinciding with the position of Zic2-expressing iRGCs (**Figure 6A** and data not shown), which is consistent with its upregulation in Zic2-electroporated neurons. By immunostaining, we also confirmed the downregulation of *Apc2* in the growth cone of Zic2-expressing neurons (**Figure 6B**), consistent with previous studies suggesting that Apc regulates axon behavior and is required for midline crossing (Arbeille and Bashaw, 2018; Purro et al., 2008). Intriguingly, also consistent with the known role of Apc2 as an integrant of the βcatenin degradation complex, we also observed a significant and homogeneous accumulation of βcatenin in the same terminals (**Figure 6B**). Lgr5, a G-protein-coupled-responding receptor proposed to trigger the canonical Wnt signaling pathway upon ligand binding (de Lau et al., 2011), was also upregulated by Zic2 (**Figure 5J**). These results indicate that although *Ctnnb* (the gene encoding βcatenin) transcription is not directly affected by Zic2 (**Figure S3B**), the presence of this TF indirectly regulates the levels of βcatenin and produces a uniform accumulation of this protein in the neuron. This βcatenin accumulation did not activate however the canonical pathway as electroporation of Top-RFP plasmids in Zic2 neurons did not result in RFP expression. In fact, retinas electroporated with Zic2, Top-RFP and Δ90-βcat plasmids showed a significant reduction in RFP expression compared to the control (**Figure 6D, E**), indicating that instead of inducing canonical Wnt signaling, Zic2 is able to inhibit this pathway. On the contrary, a significant inhibition of βcatenin-dependent transcription was observed (**Figure 6D, E**), consistent with our observation that En2, a TF involved in the nuclear translocation of βcatenin (Lin et al., 2018), is also downregulated upon Zic2 expression (**Figure S2**).

**Figure 6.**
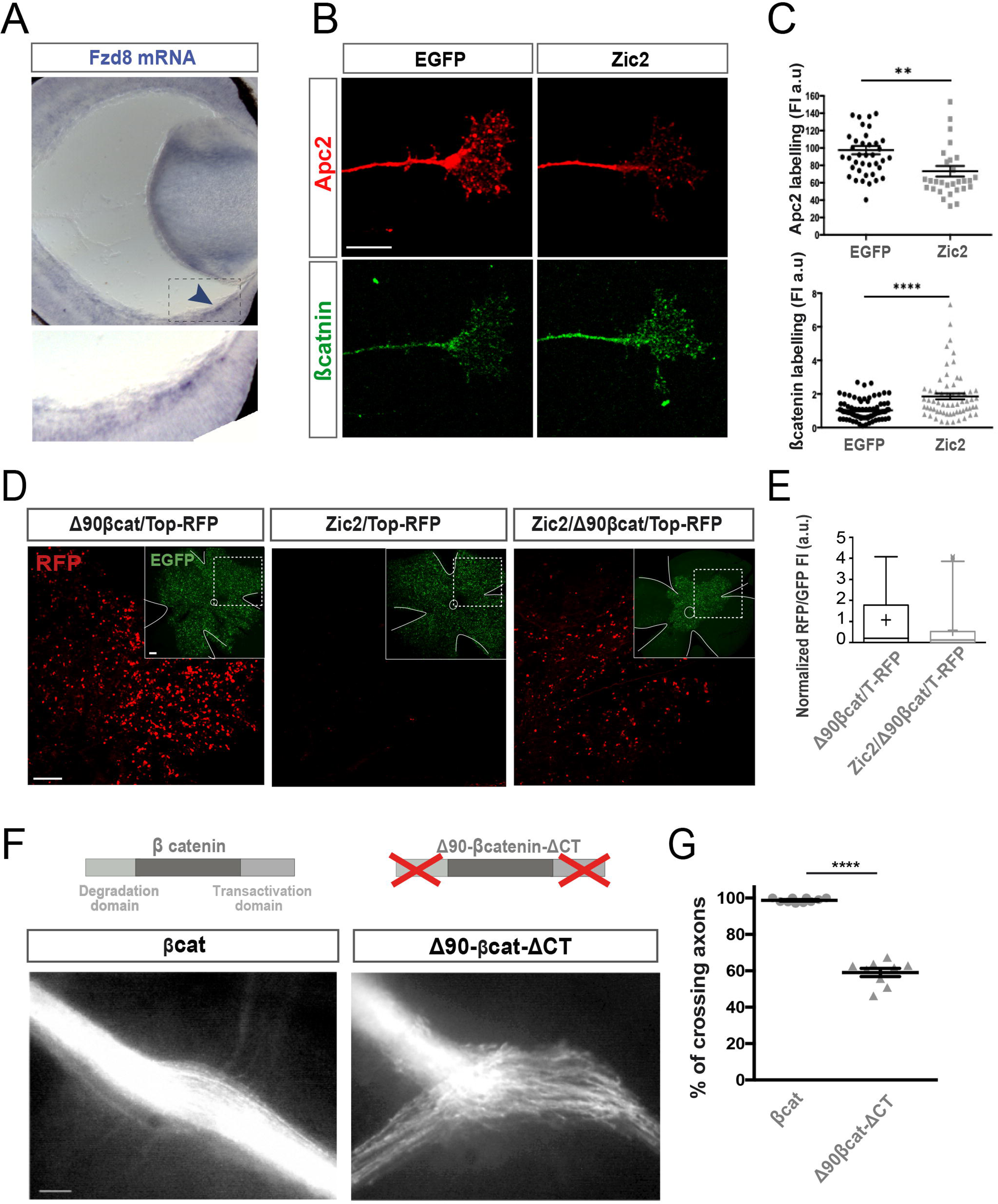
Zic2 causes accumulation of βcatenin in RGCs to disrupt midline crossing. **(A)** *In situ* hybridization for the Wnt receptor Fzd8 in a coronal section of an E16.5 mouse retina (blue arrow and higher magnification of the square are at the bottom. **(B)** Immunofluorescence for Apc2 and βcatenin in the growth cone of axons growing from explants electroporated with EGFP-or Zic2-encoding plasmids. Zic2-expressing retinal ganglion cell axons show reduced levels of the Apc2 protein and concomitant increased levels of βcatenin. Scale bar: 10μM. **(C)** Quantification of immunofluorescence intensity for Apc2 (upper graph) (n_eGFP_=41, n_Zic2_=30, p=0.0022) and βcatenin (lower graph) (n_eGFP_=81, n_Zic2_=62, p<0.0001). **(D)** Representative immunohistochemistry for RFP and EGFP in wholemount E16.5 retinas electroporated at E13.5 with plasmids encoding Δ90-βCatenin plus Top-RFP and EGFP, Zic2 plus Top-RFP and EGFP, or Δ90-βCatenin/Top-RFP/EGFP and Zic2. Images at the right corners depict targeted/EGFP cells in whole mounted retina. RFP staining shows activation of the canonical Wnt signaling. Despite the higher levels of βcatenin observed in Zic2-expressing cells versus the controls, the canonical Wnt signaling is not activated. Instead, Zic2 is able to dampen the canonical Wnt signaling in neurons that overexpress a stabilized form of βcatenin (Δ90βCat). Scale bar:100μm. **(E)** Box whisker plot shows RFP fluorescence in wholemount electroporated retinas normalized to control. Δ90-βCatenin/Top-RFP/EGFP data: n=675, Mean= 1.076 SE= 0.06 25% percentile= 0.0 Median= 0.21, 75% percentile= 1.772. Δ90-βCatenin/Top-RFP/EGFP/Zic2 data: n=649, Mean= 0.53, SE= 0.04, 25% percentile=0.00 Median= 0.13, 75% percentile= 0.43. (Two-tailed unpaired t-test ****p<0.0001). Results from three biological replicates. Results show mean ± SEM. **(F)** Optic chiasms from E16.5 embryos electroporated with plasmids encoding for βcatenin or the stabilized form of this protein Δ90-βcatenin-ΔCT (transcriptionally inactive). Scale bar: 100μM **(G)** Quantification showing the percentage of contralaterally projecting axons normalized to the total number of EGFP axons at the chiasm for each condition (βcat n=10; Δ90-βcat-ΔCT n=9) (Two-tailed unpaired t-test; **** p<0.0001. Results show mean ± SEM.

Downregulation of βcatenin provokes axon midline stalling and Zic2 induces uniform accumulation of βcatenin. These results inspired the idea that Wnt5a-induced polarized accumulation of βcatenin in the axon terminal is essential for crossing and spoiling this polarization would disturb crossing. To address this hypothesis, we analyzed E16.5 embryos electroporated at E13.5 with plasmids encoding either wildtype (wt) βcatenin or a variant form of this protein that is resistant to Apc2-mediated degradation and also lacks the transactivation domain (ΔCT-βcat). The chiasm of embryos electroporated with wt βcatenin-plasmids were undistinguishable from the controls electroporated with EGFP encoding plasmids alone. However, forced unpolarized accumulation of βcatenin achieved by the expression of this nondegradable truncated protein lead to an almost random projection phenotype with axons choosing either the ipsi or contralateral route (**Figure 6F-G**).

### Asymmetric phosphorylation of βcatenin facilitates axon steering

Previous reports have shown that the tyrosine kinase receptor EphB1, a direct target for Zic2, is necessary for the ipsilateral projection to form. However, electroporation of EphB1 in RGCs was known to induce midline avoidance with much less efficiency than electroporation of Zic2 (García-Frigola et al., 2008; Petros et al., 2009). This, already pointed to the existence of mechanisms, other than EphB/ephrinB signaling, involved in axon midline avoidance. However, such mechanisms had remained unknown until now. Together with our results showing that disrupting polarized accumulation of βcatenin produces a random projection at the midline, these observations suggested that inducing EphB1 expression in cRGCs at the same time that abolishing the capacity of the axon to respond to Wnt5a should increase the number of axons steering at the midline compared to when EphB1 is expressed alone. To test this idea, we electroporated EphB1- and *Ctnnb shRNA-*encoding plasmids at E13.5 and analyzed the optic chiasms of electroporated embryos two days later. As expected, blocking the capacity of the axons to respond to Wnt5a by reducing the levels of βcatenin and concomitantly inducing the expression of EphB1, almost doubled the number of axons changing their laterality compared to when only EphB1-plasmids were electroporated (**Figure 7A-C**).

**Figure 7.**
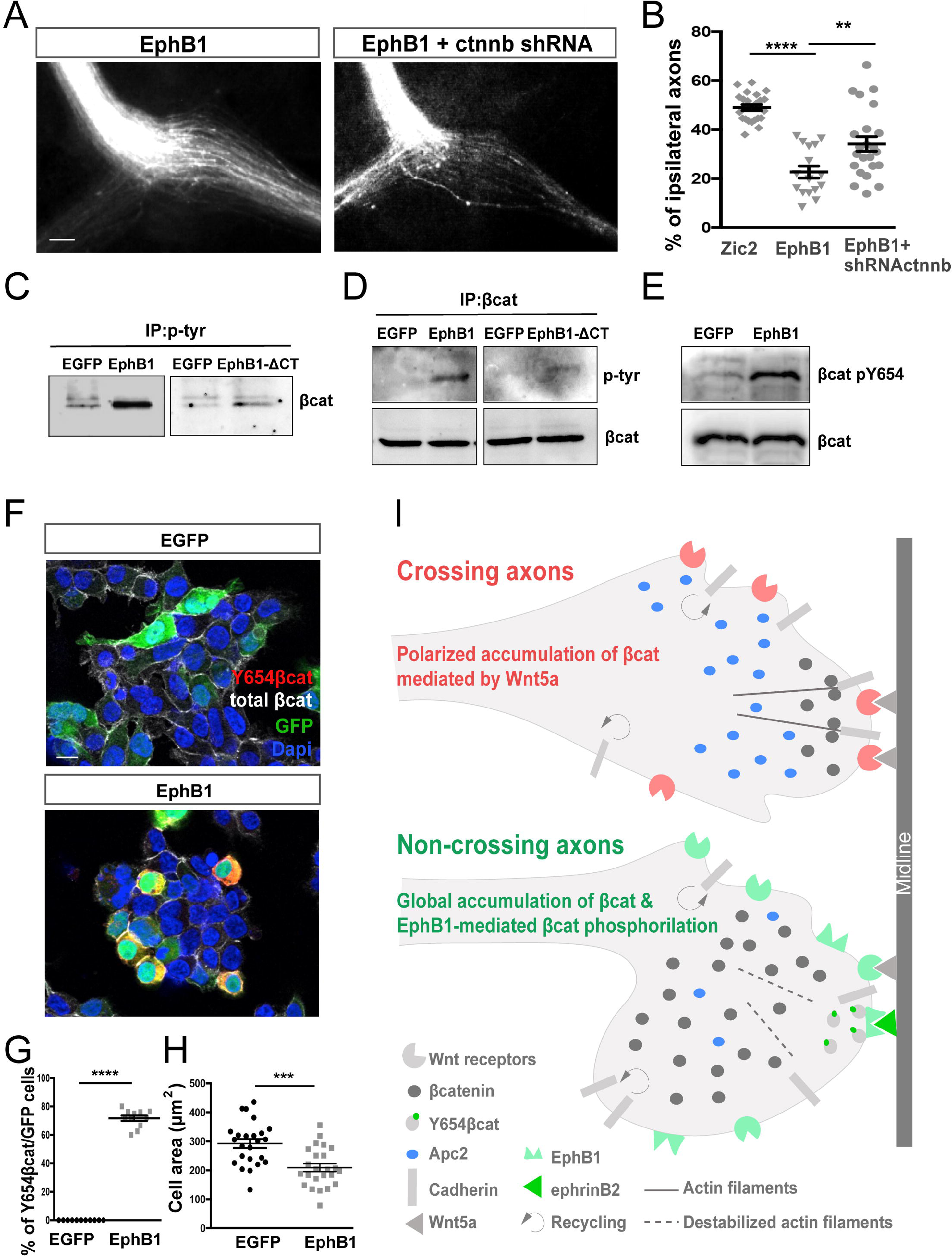
The knockdown of βcatenin facilitates EphB1-mediated axon steering. **(A)** Optic chiasms from E16.5 embryos electroporated at E13.5 with plasmids encoding EGFP plus EphB1 alone or together with plasmids bearing shRNA against βcatenin (ctnnb shRNA). Scale bar: 100μM. **(B)** Quantification showing the percentage of ipsilaterally projecting axons in each condition normalized to the total number of EGFP axons at the chiasm (Zic2 n=23; EphB1 n=16; EphB1/ctnnb shRNA n=28)(Two-tailed unpaired t-test; **p<0.01, ****p<0.0001). Results show mean ± SEM. **(C)** Detection of βcatenin in the immunoprecipitation of tyrosine-phosphorylated proteins from GFP-, EphB1- and EphB1-ΔCT-transfected cells. **(D)** Detection of tyrosine-phosphorylation in βcatenin immunoprecipitated from GFP-, EphB1- and EphB1-ΔCT-transfected cells. **(E)** Detection of Y654βcatenin in protein extracts from GFP- and EphB1-transfected cells. **(F)** Immunohistochemistry using antibodies to Y654 catenin and total βcatenin in Hek293 cells transfected with EGFP or μM. **(G)** Percentage of phosphoY654βcatenin cells on EGFP-expressing cells. Eleven ROIs were quantified from three independent experiments for each condition. Two-tailed Mann Whitney-test **** 0<0.0001. Results show mean ± SEM. **(H)** Quantification of the area occupied by individual cells transfected with EGFP alone or EphB1 plus EGFP-encoding plasmids (EphB1 n=24; EphB1/ctnnb shRNA n=24). Two-tailed unpaired t-test ***p<0.001. Results show mean ± SEM. **(I)** Working Model. As the axon grows in the absence of Wnt5a, cadherins are constantly being recycled from the plasma membrane. When contralaterally projecting axons reach the midline, the binding of Fz receptors to Wnt5a triggers the local accumulation of βcatenin linking cadherins and actin microtubules to promote cytoskeleton stabilization and facilitating midline crossing. In ipsilaterally projecting neurons, Zic2 activates a different set of Wnt receptors and other intracellular Wnt proteins at the time that it downregulates Apc2 to favor accumulation of βcatenin. Asymmetric phosphorylation of βcatenin mediated by activation of EphB1 (also induced by Zic2) through ephrinB2 binding, breaks cadherin/actin complexes and induces local microtubule destabilization.

These results suggested that ipsilateral neurons need to silence the positive response to Wnt5a to prevent crossing at the same time that inducing asymmetric cytoskeleton destabilization upon activation of EphB1 by ephrinB2 binding to promote steering. Although there is no evidence showing interactions between βcatenin and Eph receptors, crosstalk between these membrane proteins and Wnt signaling has been suggested in oncogenic contexts (Clevers and Batlle, 2006). βcatenin links membrane integrated cadherins to actin filaments and promotes cytoskeleton stabilization (Kim and Lee, 2001; Piedra et al., 2001; Roura et al., 1999). Both the levels and subcellular localization of βcatenin are tightly regulated by phosphorylation (Valenta et al., 2012). Taking all these observations into account, we considered the idea that EphB1 phosphorylates βcatenin to facilitate asymmetric cytoskeleton disassembly in ipsilaterally projecting neurons.

To tackle this hypothesis, we activated the EphB1 receptor by overexpression (Wimmer-Kleikamp et al., 2004) in HEK293 cells and performed immunoprecipitation (IP) assays using antibodies against βcatenin and phospho-tyrosine. We also transfected the cells with an EGFP-encoding plasmid or with plasmids containing a truncated version of the EphB1 receptor lacking the kinase domain (EphB1-ΔCT) as a control. We detected a large enrichment of βcatenin in the pool of immunoprecipitated tyrosine-phosphorylated proteins obtained from EphB1-transfected cells compared to GFP- and EphB1-ΔCT-transfected cells (**Figure 7C**). Consistently, we detected tyrosine-phosphorylation at a band that corresponds to the molecular weight of βcatenin in IPs using an antibody against βcatenin. This phosphoTyr-band was stronger in extracts from EphB1-transfected cells than from GFP- and EphB1-ΔCT-transfected cells (**Figure 7D**). Both experiments indicate that βcatenin is phosphorylated at a tyrosine residue/s by EphB1. Furthermore, mass spectrophotometry analysis of the immunoprecipitated product revealed that βcatenin phosphorylation occurs at the tyrosine residue 654 (Y654) upon EphB1 activation (**Figure 7E**).

Y654-βcatenin has a reduced affinity for the cadherins integrated in the cell membrane (Kim and Lee, 2001) and consistent with this, we detected Y654-βcatenin by immunofluorescence at the cytoplasm rather than in the plasma membrane and totally excluded from the nuclei of cells transfected with EphB1 plasmids. In addition, Y654-βcatenin cells exhibited smaller areas and adopted rounded shapes compared to control cells that occupied an extended area and exhibited more polygonal silhouettes (**Figure 7F-H**). These results revealed an increase in cell detachment induced by EphB1-mediated phosphorylation of βcatenin and support a model in which, EphB1 asymmetrically phosphorylates βcatenin in the growth cone upon activation by ephrinB2 at the midline to induce polarized cytoskeleton destabilization that allow axon steering.

## DISCUSSION

### An alternative βcatenin dependent but not-canonical Wnt pathway acts in axon guidance

The participation and the mode of action of the Wnt signaling pathways in axon guidance has been a controversial issue for years (Avilés and Stoeckli, 2016; Bovolenta et al., 2006; Domanitskaya et al., 2010; Lyuksyutova et al., 2003; Wolf et al., 2008). Our results reconcile all the previous observations about the participation of the canonical versus the non-canonical Wnt pathway in axon navigation, by revealing a third via that regulates crossing and it is independent on βcatenin-mediated transcription but relays on its local accumulation. This form of signal transduction is remarkably different to the canonical pathway because the changes in βcatenin stability are restricted to the axon terminal compartment and never reaches the cell nucleus nor trigger transcription. However, because it involves βcatenin dynamics is not the classical PCP pathway either.

In contralaterally projecting axons, Wnt5a binding induces polarized accumulation of βcatenin, which in turn leads to cytoskeleton stabilization at the tip of the growth cone and promotes midline crossing. In contrast, the presence of the transcription factor Zic2 in ipsilaterally projecting neurons, induces a gene program that silences this positive response to Wnt5a and triggers instead EphB1-dependent phosphorylation of βcatenin that allows cytoskeleton destabilization (**Figure 7I**).

Interestingly, our genomic analysis also revealed that from those Wnt-related genes showing Zic2 occupancy, only a subset is differentially expressed after Zic2 induction (e.g., *Fz1*, *Fz8*, *Lrg5*, Apc2, Sox4 or *Diversin*/*Annkr6*). We did not find Zic2-peaks in genes specifically associated with the PCP pathway, such as VanGogh/Strabismus (*Vang*/*Stbm*), Flamingo (Fmi), Prickle (*Pk*), JNK, c-jun, or AP-1 except for *Diversin*/*Ankrd6*, a protein that recruits CKI-epsilon to the βcatenin degradation complex and has been described as a molecular switch between the canonical and the non-canonical pathway (Jones et al., 2014; Schwarz-Romond et al., 2002).

Together, these findings clarify many aspects of the long-standing debate about Wnt signaling pathways in axon pathfinding. First, this work shows that Wnt5a elicits attractive or repulsive responses in the same type of neurons depending on a particular transcriptional program that controls the expression of a set of receptors and specific intracellular proteins. Moreover, they reveal that axon repulsion cannot be achieved only through the action of repulsive receptors since ipsilateral axons need to silence attractive mechanisms for effective steering. Finally, they also clarify that Wnts do not activate the canonical or PCP pathways in axon guidance processes but complex mechanisms based on βcatenin dynamics at the growth cone.

### Zic2-silencing of the alternative Wnt pathway likely operates in other contexts

Our study does not only reveal a new branch of the Wnt signaling pathway, it also identifies the TF responsible for the two-way switch in signal transduction. We propose that a similar switch in the response to Wnt ligands may operate in other tissues and contexts where Wnt signaling and Zic2 coexist beyond axon guidance. For instance, Zic proteins are expressed in several populations of migrating neuroblasts in the forebrain, hindbrain and neural tube as well as in neural stem cells in the adult individual (Herrera, 2018; Inoue et al., 2007; Merkle et al., 2014; Murillo et al., 2015; TeSlaa et al., 2013a; Tiveron et al., 2017). In all these contexts, cells expressing Zic TFs are in contact with Wnt proteins (Herrera, 2018) and delaminate from a neuroepithelium undergoing shape rearrangements before migrating through stereotyped paths. Ipsilaterally projecting neurons, or at least a significant number of them, also originate from an epithelium-like area known as the ciliary marginal zone (Fernández-Nogales et al., 2019; Marcucci et al., 2016) and the cell bodies of these cells are exposed to Wnt ligands secreted from this peripheral retinal area (Denayer et al., 2008; Meyers et al., 2012). In these cells, Zic2 likely plays a two-fold function regarding Wnt signaling. In addition to regulate axon guidance as we show here, it would block the canonical pathway to prevent proliferation, guaranteeing that the recently differentiated neurons do not reenter the cell cycle. Consistent with this idea, it has been shown that activation of Wnt signaling alters the number of ipsilaterally projecting neurons potentially by keeping them in the cell cycle (Iwai-Takekoshi et al., 2018). Furthermore, experiments in zebrafish and human cell lines suggest that Zic2 blocks the canonical Wnt signaling pathway (Pourebrahim et al., 2011), a result that we have also confirmed here (**Figure 3E, F**) and that could occur in particular contexts concomitantly with its role in regulating the alternative Wnt pathway.

Zic2 mutations in mice and humans cause holoprosencephaly type IV, anencephaly and spina bifida (Brown et al., 1998; Nagai et al., 2000) likely as a consequence of defects in early developmental stages such as gastrulation and/or neurulation (Houtmeyers et al., 2016; TeSlaa et al., 2013b). In addition, accumulating evidence shows that this TF is upregulated in many different types of cancer (for a recent review see Houtmeyers et al., 2018). However, the molecular mechanisms underlying these Zic2-associated anomalies have remained largely elusive. Recent reports have also shown novel roles for Wnt signaling in the injured CNS (Garcia et al., 2018). Wnt and Eph/ephrin pathways both play key functions in all these processes. While further experiments should revisit the role of Zic2 in early development and oncogenic scenarios, our results support the idea that Zic2 may regulate the crosstalk between the Eph and Wnt pathways in many different contexts.

## ACKNOWLEDGEMENTS

We thank Y. Coca, and M. Herrera for technical assistance. We also thank P. Bovolenta and C. Mason for discussion and comments on the manuscript. The laboratory of EH is funded with the following grants: (BFU2016-77605 from the National Grant Research Program, PROMETEO Program (2016/026) from Generalitat Valenciana, (PCIN2015-192-C02-02 from ERA-Net Program) and (ERC-282329 from the European Research Council). MLC is the recipient of a FPI fellowship from the National Grant Research Program. J. L-A research is supported by grants RYC-2015-18056 and RTI2018-102260-B-100 from MICINN co-financed by ERDF. A.B. research is supported by grants SAF2017-87928-R from MICINN co-financed by ERDF, PROMETEO/2016/026 from the Generalitat Valenciana. We also acknowledge the financial support received from the “Severo Ochoa” Program for Centers of Excellence in R&D (SEV-2013-0317).

## AUTHOR CONTRIBUTIONS

C M-P designed, performed and analyzed most of the experiments. MT L-C and JP L-A have performed the computational analysis of RNA-seq and ChIP-seq data. DB performed *in situ* hybridizations and took care of the mice colonies. L C-D and A G-G performed some immunostainings. EH wrote the original draft and designed, conceived and supervised the study. AB revised subsequent versions of the manuscript and helped with experimental design and revisited critically the manuscript for important intellectual content.

## MATERIAL AND METHODS

### Mice

All experiments were performed using embryos from C57/DBA F1 hybrids. Mice were kept in a timed pregnancy breeding colony at the Instituto de Neurociencias (IN). The animal protocols were approved by the IN Animal Care and Use Committee and met European and Spanish regulations.

### Cell culture, plasmids and luciferase assay

HEK293 cells (ATCC® CRL-15736™) were cultured according to standard conditions. For luciferase assays, cells were transfected using Lipofectamine 2000 (Invitrogen) according to manufacturer’ guidelines. Complementary primers containing Zic2 binding motifs A or B were cloned in pGL3-basic (See Supplementary methods for sequences) and co-transfected with or without CAG-Zic2 plasmid in conjunction with a thymidine-kinase (TK) promoter-β-Gal. Cell lysates were harvested the day after and luciferase and β-galactosidase activities were measured using Luciferase Reporter and Beta-Glo Assay Systems (Promega) following the manufacturer’s guidelines. Luciferase activity of all transfections was normalized to β-Gal activity.

### In utero electroporation and quantification

Time-pregnant mouse females were anesthetized with a classic small animal anesthesia (isoflurane) system (WPI, USA). In utero electroporation was performed as described in García-Frigola et al. (2008). Plasmids with pCAG-Zic2, pCAG-Δ90-βCatenin-GFP (Addgene 26645) pCAG-Full-βcatenin-GFP, pCAG-Δ90-βCatenin-ΔCT-GFP, pCAG-EphB1, Top-RFP were injected at 1μg/μl and pSilencer-shCtnnb1 and pSilencer-control at 2 μg/μl. pCAG-GFP plasmids were co-injected at 0.5μg/μl. pCAG-Δ90-βCatenin-ΔCT-GFP and pCAG-Full βCatenin-GFP were obtained from manipulation of the CAG-Δ90-βCatenin-GFP (See Supplementary Methods for details). Fiji software was used to quantify ipsi and ipsilateral projections. Briefly, mean fluorescent intensity from three linear ROIs drawn perpendicularly on each optic tract (OTF) was measured and background from a very proximal area was rested to each measure. The % of ipsilateral projection was determined for each individual embryo by applying the following formula: % Ipsis= (iOTF x100)/(iOTF+cOTF). Statistical analyses were performed with GraphPad Prism 6.0 (GraphPad software, Inc, La Jolla, CA).

### Retinal explant cultures and immunohistochemistry

Retinal explants from E14.5 WT or electroporated embryos were plated on glass bottom microwell dishes (MatTek) coated with poly-L-lysine 0.01% and laminin (20μg/ml). Culture media was DMEM:F12/Neurobasal media (Gibco TM) with 0.4% of methylcellulose with growth supplements N-2 and B-27 and antibiotics Pen/Strep. For axonal growth analysis recombinant Wnt5a at 100ng/ml or Wnt5a reconstitution buffer as a vehicle was added to the medium. For acute responses, retinal explants were exposed to recombinant Wnt5a at 200ng/ml for 1 hour.

Immunohistochemistry was performed on retinal explants fixed with pre-warmed 4% PFA in PBS at 37°C for 20 min. Explants were permeabilized with 0.025% Triton X-100, blocked with horse serum and incubated with the specified antibody at 4°C overnight. Fluorescence microscopy was performed using a Leica confocal microscope SPEII. Area and fluorescence intensities at the growth cones were quantified by Fiji software using maximum intensity z-projection.

### In situ hybridization

Heads of embryos from E13.5 to E17.5 were dissected in cold PBS 1X and fixed in 4% PFA overnight. Coronal vibrosections (optic chiasms) or criosections (retinas) were obtained and *in situ* hybridization with specific antisense riboprobes for different Wnts and Wnt receptors (gift of Prof. P. Bovolenta). Images were captured with a Leica DM2500 equipped with a Leica DFC7000T camera and Leica Application Suite. Version 4.10.0 Software.

### RNA-seq, ChIP-seq and bioinformatic analyses

RNA-seq: RGCs from E13 electroporated retinas were isolated 36 h post-electroporation by cell sorting. Isolated retinas were enzymatically dissociated in a mixture of collagenase/Trypsin and BSA for 20 minutes at 37°C followed by mechanical dissociation. Single cell suspension was filtered and resuspended in cold HBSS medium supplemented with 20% FBS. After cell sorting, cells were centrifuged and frozen. Total RNA extraction was performed with Arcturus® Picopure® RNA isolation Kit (ThermoFisher Scientific). The samples were sequenced according to the manufacturer instructions in a HiSeq Sequencing v4 Chemistry (Illumina, Inc). Briefly, RNA-seq reads were mapped to the mouse genome (Mus_musculus.GRCm.38.83) using STAR (v2.5.0c) (Dobin et al., 2013). Quality control of the raw data was performed with FastQC (http://www.bioinformatics.babraham.ac.uk/projects/fastqc/). Library sizes were between 41 and 75 million single reads. Samtools (v1.3.1) were used to handle BAM files (Li et al., 2009a). To retrieve differentially expressed genes (DEG), mapped reads were counted with HTSeq v0.6.1 (Anders et al., 2015) with the following parameters: -s reverse –i gene_id and with a gtf file reference of GRCm38.83. Read count tables were analyzed using DeSeq2 v1.10.0 (Love et al., 2014). Analysis and preprocessing of data were performed with custom scripts using R (https://cran.r-project.org/) (v3.4.3“Kite-Eating Tree”) statistical computing and graphics, and bioconductor v3.2 (BiocInstaller 1.20.3) (Huber et al., 2015). Genes were considered differentially expressed at Benjamini–Hochberg (BH) adjusted *p*-value□<□0.1 Significantly upregulated and downregulated genes were visualized with IGV (v2.3.72) (Thorvaldsdóttir et al., 2013). GO enrichment analyses were performed using the platform Panther (Mi et al., 2019), with Fisheŕs exact test and with the Padj correction, obtaining the top terms using the filters by ratio enrichment > 2, number of GO family group genes between 3 and 2000, number of enrichment genes >3, and Padj < 0.1. The gene set enrichment analysis was obtained with GSEA v3.0 (Subramanian et al., 2005).

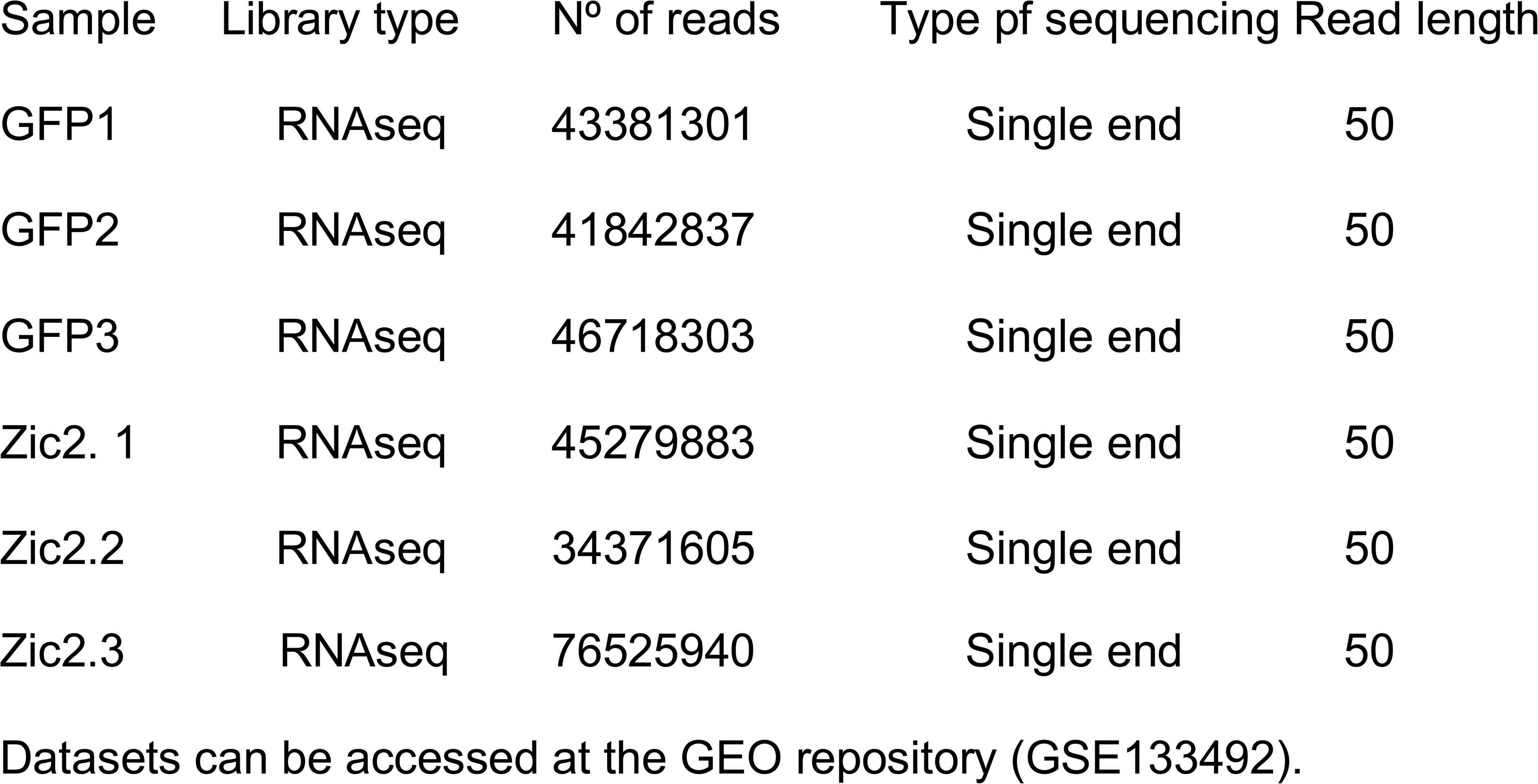

ChIP-seq: Whole retinas or spinal cords were dissected from embryos at E16, chopped, crosslinked with 1% PFA at room temperature for 20 minutes and quenched with glycine. Then, chromatin was sheared using a bioruptor sonicator (Diagenode) and immunoprecipitated as described in STAR Methods. Breafly, sheared chromatin was diluted and incubated overnight at 4°C with the specific antibody against mouse Zic2 (Millipore, AB15392) or IgG isotype control pre-immune serum (Abcam, ab27478). Immunoprecipitated chromatin was purified with Qiaquick PCR Purification Kit (Quiagen). To reduce biological variability, for each independent sample, chromatin was obtained by pooling tissue from several mice (retina, 176 mice; spinal cord, 68 mice). NGS Libraries were prepared from purified DNA from anti-zic2 ChIP (Zic2, IgG pre-immune serum) and input samples according to manufacturer instructions (Illumina). Libraries were single-end sequenced in HiSeq2000 apparatus using Flow Cell v3 chemistry (read length: 1 x 50 bp). Information on library preparation method, size of the libraries, and mapping to reference genome can be found at the end of this section. Quality control of the raw data was performed with FastQC (http://www.bioinformatics.babraham.ac.uk/projects/fastqc/). Libraries aligned to the mouse genome (mm9, NCBI assembly M37) using Burrous-Wheeler Alignment Tools (BWA) (v0.5.9) (Li and Durbin, 2010) and further processed using samtools (v0.1.17) (Li et al., 2009b) and bedtools (v2.19.0) (Quina, 2005). Additional quality control analyses on aligned sequences from of ChIP and input samples were performed using the R package Repitools (Statham et al., 2010). Peak calling for Zic2 ChIP-seq was (Li and Durbin, 2010) performed using MACS2 (v1.4.2) (Zhang et al., 2008) with the following parameters: -p 1.00e-05 –bw 300 -m 10,30. Read counts on aligned bam files were obtained using featureCounts (Rsubread v1.12.6) (Liao et al., 2014). Further data processing was performed with custom scripts in the R programming language (https://cran.r-project.org/). *De novo* DNA motif discovery at Zic2-bound regions was performed using Homer software (Heinz et al., 2010) and MEME suite (Bailey et al., 2009), in an area of 151 bp (average genomic fragment length) around each peak summit. GO enrichment analyses were performed using the platform Panther, with Fisheŕs exact test and with the Benjamini-Hochberg False Discovery Rate for multiple test correction for the nominal *p* values. GO top terms were obtained using the following filters: Padj < 0.1, ratio enrichment > 2, number of genes in enriched term >3, number of GO family group genes between 3 and 2000. Heatmaps of ChIP-seq read density and aggregate density plots were performed using SeqMiner (v1.3) (Ye et al., 2011) and R custom scripts. ChIP-seq peaks were annotated using ChIPpeakAnno (Zhu et al., 2010). ChIP-seq tracks were visualized using IGV (v2.3.72) (Robinson et al., 2011).

| Sample | Library type | N° of reads | Type pf sequencing | Read length |
| --- | --- | --- | --- | --- |
| Input_dSCN | ChIP DNA | 46419753 | Single end | 50 |
| Zic2_dSCN | ChIP DNA | 31047920 | Single end | 50 |
| Input_RGCs | ChIP DNA | 94319654 | Single end | 50 |
| Zic2_RGCs | ChIP DNA | 64787935 | Single end | 50 |

Datasets can be accessed at the GEO repository (GSE133492).

### Immunoprecipitation, mass spectrometry and Western-Blot

HEK293 cells were transfected with pCAG-FullβCatenin-GFP or pCAG-FullβCatenin-GFP/pCAG-EphB1 and isolated by cell sorting 48 hours post-transfection. Total protein was extracted in RIPA buffer with protease and phosphatase inhibitors. To obtain a phosphotyrosine-enriched βcatenin sample a tandem enrichment strategy was used. First immunoprecipitation was performed with an anti-phosphotyrosine antibody, protein-G bounded proteins were eluted with phenyl-phosphate (100mM) and then immunprecipitated with anti-βCatenin antibody. Samples were analyzed by mass spectrometry. Protein Identification by LC/MS/MS (LTQ-Orbitrap-Velos) was carried out in the ‘CBMSO Protein Chemistry Facility, that belongs to ProteoRed, PRB2-ISCIII, supported by grant PT13/0001.

### Data Collection and Analysis

Data collection has been performed using the following equipment: FACS Aria2, Illumina NovaSeq; Leica confocal microscope SPEII; Leica DM2500; Leica DFC7000T camera and CUY21SC electroporator. Data analysis has been performed using the following softwares: Flowjo v10; Fiji software maximum intensity z-projection; Leica Application Suite (LAS) Software; Image J(v2), Microsoft Excel, (v16); FastQC (v.0.11.6), RSeQC (v2.6.4), fastq-dump (v2.3.5), Python 3.7.3, R software (v3.4.4), Cutadapt (v1.18), Samtools (v1.9), Bedtools (v2.26.0), Rsubreads (v1.26.0), STAR(v2.5.0a), HTSeq (v0.7.2), DESeq2 (v1.10.0), Bowtie2 (v2.2.6), MACS2 (v2.1.0), ChIPpeakAnno (v.3.18), biomaRt (v.2.39), PANTHER (v14), IGV (v2.3.92), HOMER (v4.8), MEME Suite (v.5.0.3); GraphPad Prism Software (Version 7.0) San Diego. Statistical analysis was performed as indicated in Figure Legends. Sample size was estimated according to data variance and correlation between biological replicates and distance to control condition. Each analysis was done in accordance to the sample size, sequencing depth and conditions. Reproducibility across replicate was very high. The measures applied to evaluate replicates were correlation among samples, variance, Euclidean classification, principal component analysis (PCA) and statistical tests that revealed not significant differences within the group. In the analyses in which a replicate was not available, analyses were supported by the use of internal controls and the sample was normalized and compared with samples from related conditions in which replicates were available (i.e., ANOVA, Desq2).

