## Supplementary figures and images for "Zic2 abrogates an alternative Wnt signaling pathway to convert axon attraction into repulsion"

### Figure 1

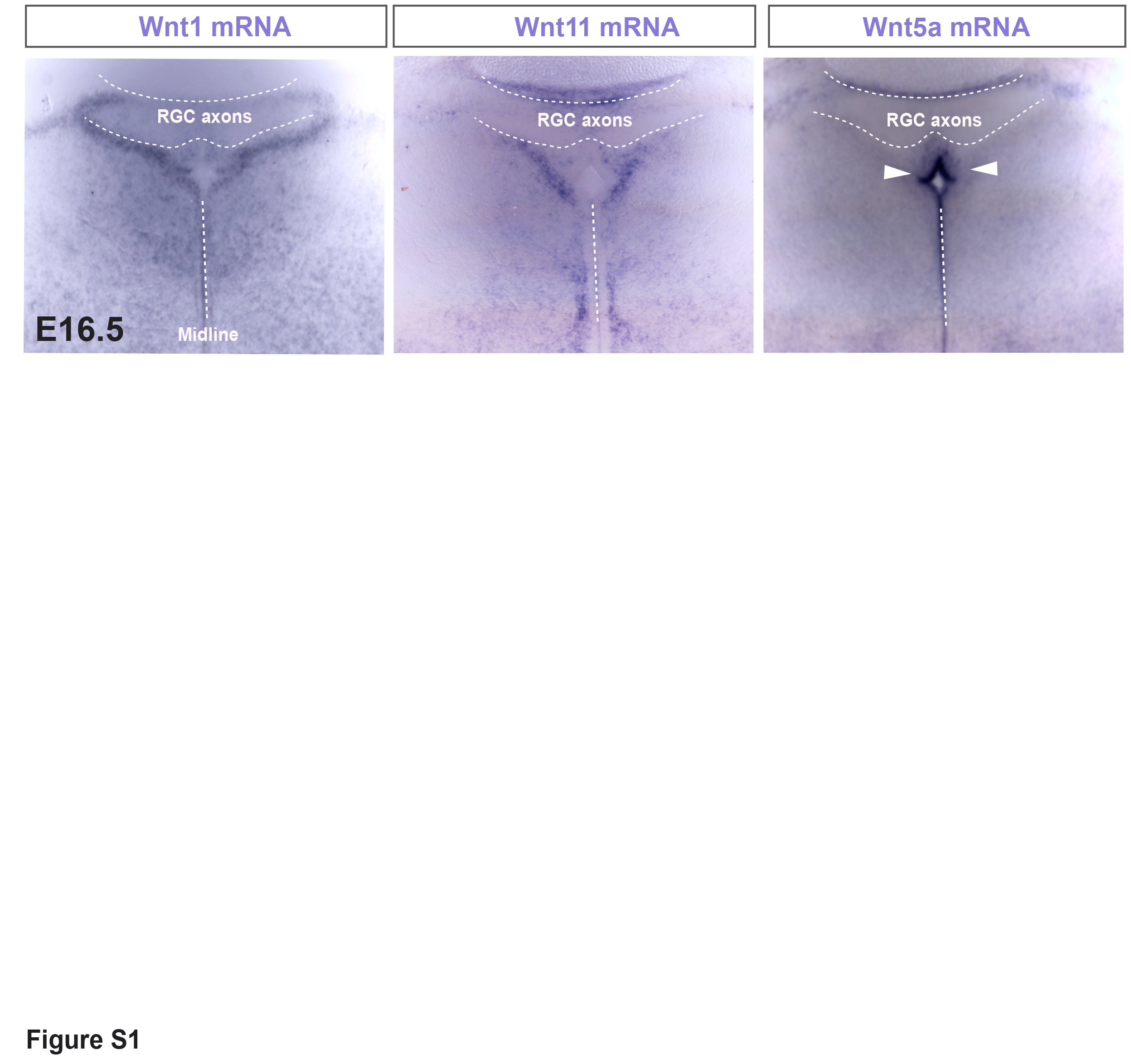

### Figure 4

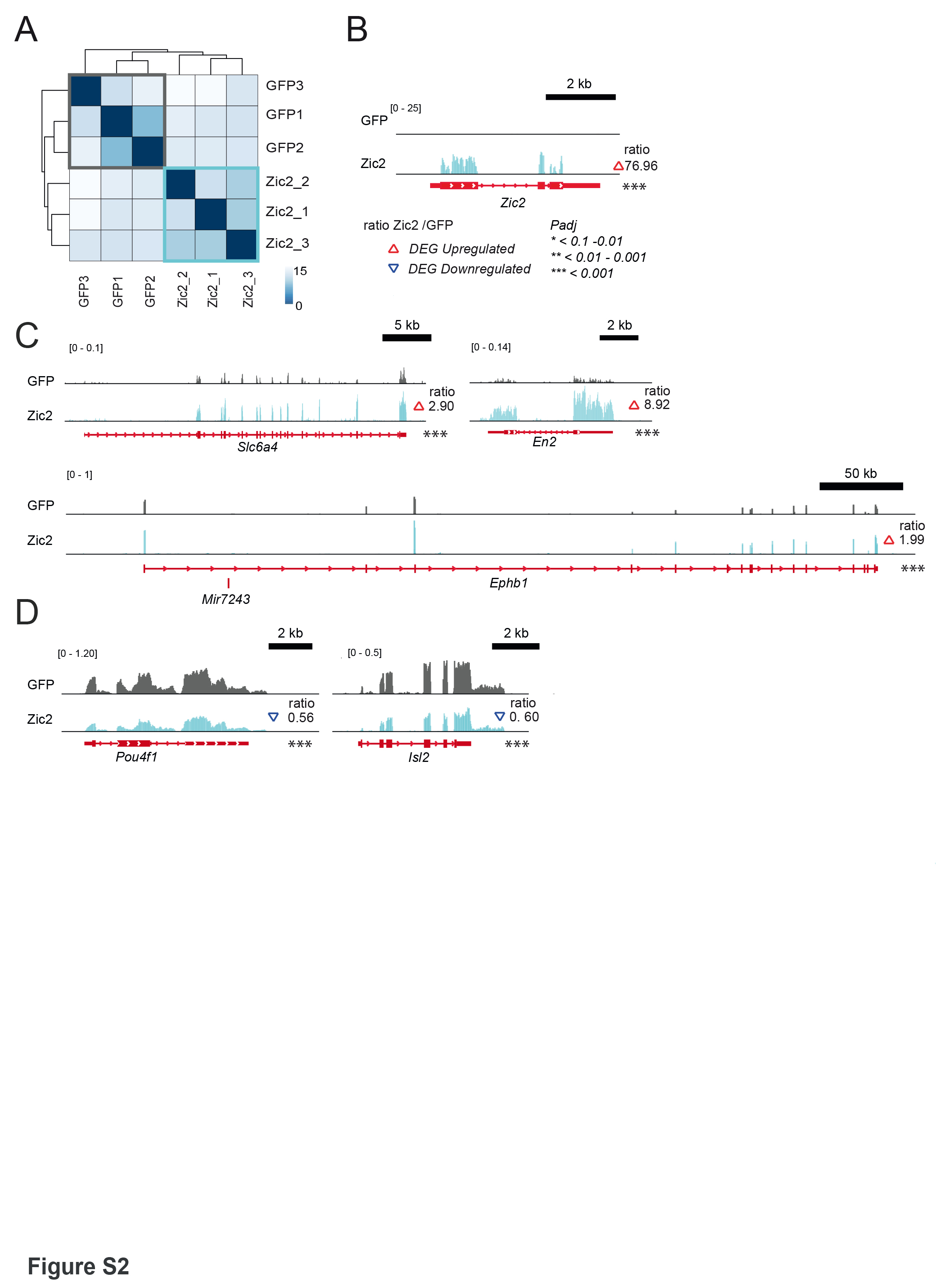

### Figure 5

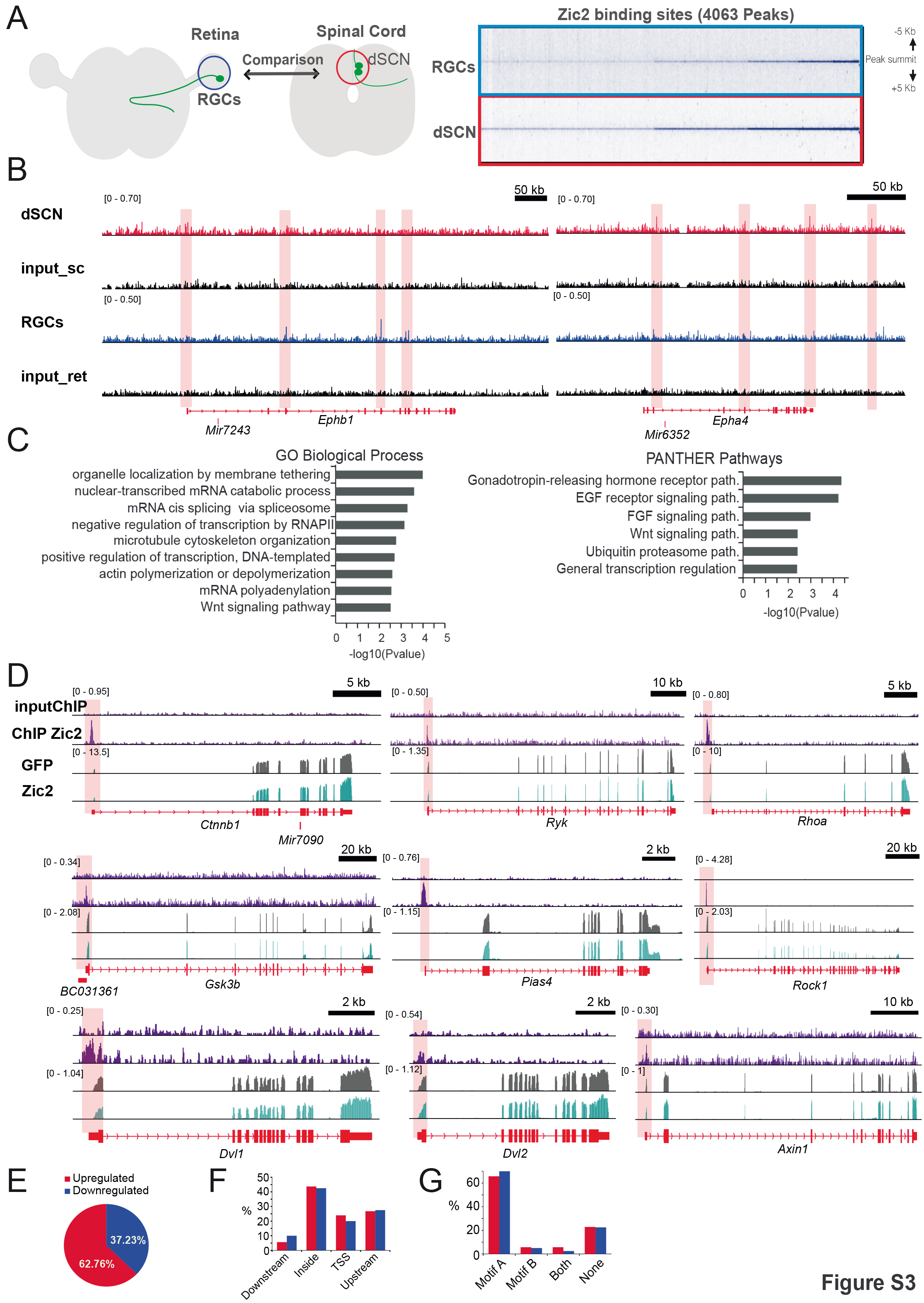
