## Supplemental Methods for "Zic2 abrogates an alternative Wnt signaling pathway to convert axon attraction into repulsion"

**LEAD CONTACT AND MATERIALS AVAILABILITY**

**Materials Availability Statement**

Plasmids generated in this study are available upon request.

**EXPERIMENTAL MODEL AND SUBJECT DETAILS**

**Cell lines**

Cell line from human embryonic kidney 293 (HEK293) was obtained from ATCC repository. General maintenance was in DMEM + GlutaMAX™ medium (Gibco ref.31966-021) supplemented with 10% fetal bovine serum and Penicillin (100 U/ml) /Streptomycin (100μg/ml). Growth conditions were 37ºC and 5% CO2. Cells were passed when they reached >80% confluency.

**Retinal Explants**

Mouse retinal explants were plated on poly-L-Lysine and Laminin and were grown in DMEM/F12 medium + 0.4% methylcellulose supplemented with N2/B-27 and Penicillin (100U/ml) /Streptomicin (100μg/ml) at 37ºC and 5% CO2.

**Animals for *In utero* electroporation**

Mice were kept in a timed pregnancy breeding colony at the Instituto de Neurociencias (IN). All experiments were performed in embryos from C57/DBA F1 hybrids at E13. Animals were housed under conventional conditions and had ad libitum access to food and water. The animal protocols were approved by the IN Animal Care and Use Committee and met European and Spanish regulations.

**METHOD DETAILS**

***Cell culture and immunocytochemistry***

HEK293 cells (ATCC® CRL-15736^TM^) were cultured according to standard conditions. For immunohistochemistry, cells were transfected using Lipofectamine 2000 (Invitrogen) according to manufacturer’ guidelines. The day after, cells were detached with 0,025% trypsin and divided in two coverslips treated with 0,01% poly-L-lysin. After 24 hours (48h post-transfection) the cells were fixed with 4% PFA in PBS, washed and incubated with blocking solution 0,025% Triton-X100, 5% Horse serum in PBS and incubated with the specific antibodies (mouse anti-βcatenin 1/500, mouse anti-p-Tyr 1/500 (ref. 05-321, Sigma Aldrich), rabbit anti-phospho-βCatenin-Y654 1/250 (ref.ab59430, Abcam), chicken anti-GFP 1/3000) and DAPI to visualize nuclei.

***DNA motifs cloning and Luciferase assay***

Motif A and B containing complementary primers were purchased from Sigma-Aldrich. Motif A: forward 5’-CGAATTC**CCTGCGCATGCGCAGTG**C-3’, reverse 5’-TCGAG**CACTGCGCGCATGCGCAGG**GAATTCGAGCT-3’ Motif B: forward 5’-CGAATTC**ACTACAATTCCCAGCATG**C-3’	rev 5’-TCGAG**CATGCTGGGAATTGTAGT**GAATTCGAGCT-3’. For luciferase assays, Hek293 cells were seeded in 24-well plates and transfected using Lipofectamine 2000 (Invitrogen) according to manufacturer’ guidelines. Complementary primers containing Zic2 binding motifs A or B were obtained from Sigma-Aldrich. They were annealed, cloned in pGL3-basic and co-transfected with or without CAG-Zic2 plasmid in conjunction with a thymidine-kinase (TK) promoter-β-Gal. Cell lysates were harvested the day after and luciferase and β-galactosidase activities were measured using Luciferase Assay System (ref.E4720) and Beta-Glo Assay Systems (ref.E1500) (Promega) following the manufacturer's guidelines. Luciferase activity of all transfections was normalized to β-Gal activity. Data from three independent experiments were analyzed.

***In utero Electroporation and quantification.***

Sh-RNA sequence to target mouse βCatenin was obtain from Genetic Perturbation Platform (GPP) database (Broad Institute). Target sequence of mouse -βCatenin was CGTGAAATTCTTGGCTATTAC (NM_007614.2-1082s21c1). Match location at position 1082 of the coding region sequence. Oligos wearing sh-RNA sequences were cloned in plasmid pSilencer 2.1-U6 Neo (Thermo Fisher Scientific. Ref AM5764). pSilencer-shβCatenin and pSilencer-control at 2 μg/μl. In all cases the above plasmids were injected in combination with pCAG-GFP at 0.5μg/μl to visualize axonal trajectories of electroporated cells. pCAG-Δ90-βCatenin-ΔCT, pCAG-Full βCatenin were obtained by amplifying the N-terminal domain (first 90 aa) of mouse βCatenin and cloned in open reading frame into pCAG-Δ90-βCatenin.

***Retinal explant cultures and Immunohistochemistry***

For axonal growth analysis, recombinant human/mouse Wnt5a protein (R&D Systems, ref. 645-WN-010) at 100ng/ml or Wnt5a reconstitution buffer as a vehicle were added to the medium and explants were grown overnight at 37ºC and 5% CO. For acute responses, retinal explants were exposed to recombinant Wnt5a at 200ng/ml for one hour at 37ºC and 5% CO_2_.

Immunohistochemistry was performed on retinal explants fixed with pre-warmed 4% PFA in PBS at 37ºC for 20 min. Explants were permeabilized and blocked with 0.025% Triton X-100, 5% horse serum in PBS for 1 hour at RT and incubated with the specified antibody at 4ºC overnight. Anti-βCatenin (1/500, ref. 610153, BD Transduction Laboratories), anti-tuj1 (1/1000, ref.ab18207, Abcam), anti-APC2 (1/500, ref. PA5-20944 Invitrogene), anti-GFP (1/3000, ref.GFP-1020, Aves Labs Inc.). Secondary alexa-conjugated antibodies were used at 1/1000 dilution (Fisher. Mounting media for further image analysis was 10% Mowiol /24% glycerol. Fluorescence microscopy was performed using a Leica confocal microscope SPEII. Area and fluorescence intensities at the growth cones were quantified by Fiji software using maximum intensity z-projection.

***In situ hybridization***

Heads of embryos from E13 to E17 were dissected in cold PBS 1X and fixed in 4% PFA overnight. Coronal vibrosections (optic chiasms) or criosections (retinas) were obtained and *in situ hybridization* was performed according to reported methods with specific antisense riboprobes for different Wnts and Wnt receptors. Retrotranscription of linearized plasmids bearing the respective cDNAs (gift of Prof. P. Bovolenta) was performed by standard methods. Images were captured with a Leica DM2500 equipped with a Leica DFC7000T camera and Leica Application Suite (LAS) Software.

***Fluorescent activated cell sorting and mRNA-seq***

Electroporated retinas (from 4 to 10 retinas per each individual isolation) with pCAG-GFP or pCAG-Zic2/pCAG-GFP were dissected 36 hours post-electroporation and retinal ganglion cell suspension was achieved by enzymatic dissociation (0.125% Trypsin-EDTA, 1mg/ml Collagenase A, 0.2% BSA, 50 units/ml Deoxyribonuclease I (Invitrogen)) at 37ºC for 20 min. followed by gently mechanical disaggregation in 20% FBS-DMEM medium. The next procedures were performed keeping the samples at 4ºC. Single cell suspension was obtained by filtering using nylon Cell strainer 40 μM (Falcon) and cells were centrifuged at 1250 rpm for 5 min. Cellular pellets were resuspended in 20% FBS-DMEM medium plus 50 units/ml of desoxirribonuclease I to achieve a cellular density about 5000 cells per ml. GFP-expressing RGCs were isolated by fluorescent activated cell sorting using BD FACSAria™ III device (BD Bioscience) and selecting the purity parameters. A small sample of segregated cells was visualized by fluorescent microscopy to check purity and quantity. Cells were pellet and frozen at -80ºC until total RNA extraction. Total RNA extraction was performed by pooling cell pellets from different litters and experiments to obtain three biological independent replicates of similar number of cells ( ≈ 7000 cells) and using the Arcturus® PicPure® RNA Isolation Kit (Thermo Fisher Scientific, KIT0204). RNA was eluted in 12 μl of RNAse-free milli-Q water (Ambion®) and sent to Center for Genomic Regulation (CRG, Spain) for sequencing service. Quality and quantity of total RNA was assessed by bioanalyzer device.

***ChIP-seq***

In total, 68 spinal cords or 180 retinas were dissected from embryos at E16.5 from different litters and in different days, chopped, crosslinked with 1.1% PFA at room temperature for 20 minutes and quenched with glycine to a final concentration of 0.125M. The rest of the procedures were performed at 4ºC and with the presence of proteases and phosphatases inhibitors (Pierce™ phosphatase inhibitor mini tablets, Complete™ protease inhibitor Cocktail tablets, Merck).To isolate nuclei, tissues were homogenize using a glass pestle in lysis buffer (50mM HEPES-KOH pH 8.0, 1mM EDTA, 0.5 mM EGTA, 140mM NaCl, 10% glycerol, 0.5% NP-40, 0.25% Triton-X100) and incubated with this buffer in rotation for 10 min. Then, intact nuclei were centrifuged at 600 x g and resuspended in washing buffer (10 mM Tris-HCL pH8.0, 1mM EDTA, 0.5mM EGTA, 200mM NaCl) and rotated for 10 minutes. Then nuclei were collected by centrifugation at 600 x g, 10 minutes. Nuclei were lysed in SDS lysis buffer (1% SDS, 10mM EDTA, 50mM Tris pH8.0) and chromatin was sheared using Bioruptor® sonication device (24 cycles (changing cold water every 4 cycles) of 30 sec. ON/ 30 sec. OFF), quantified by nanodrop and checked by agarose electrophoresis. For each individual ChIP experiment about 10-15 micrograms of diluted chromatin (0.01% SDS, 1.1% Triton-X100, 1.2mM EDTA, 16.7mM Tris-HCl pH 8.1, 167mM NaCl) were precleared with 30 μl of Protein G-Sepharose Fast Flow (blocked with 1% BSA in Ch-IP dilution buffer) (Sigma, P3296) and incubated overnight at 4ºC with the specific antibody against mouse Zic2 at 1/400 (Millipore, AB15392) or IgG isotype control pre-immune serum at 1/400 (Abcam, ab27478) after recovery the 10% of total sample volume as input. The day after, Ch-IPs were centrifuged at maxim speed in a bench centrifuge and 30 μl of 1% BSA-Protein G-Sepharose was added for 1 hour incubation at 4ºC. Chromatin-protein complexes bound to Protein G-Sepharose were washed two times with low salt buffer (0.1 % SDS, 1% Triton X-100, 2mM EDTA, 20mM Tris-HCl pH8.0, 150mM NaCl), once with high salt buffer (0.1 % SDS, 1% Triton X-100, 2mM EDTA, 20mM Tris-HCl pH8.0, 500mM NaCl), two times with LiCl buffer (0.25mM LiCl, 1% IGEPAL-CA630, 1% NaDoc, 1mM EDTA, 10mM Tris-HCl pH 8.1) and two times in TE buffer. Then, samples were eluted two times with 100 μl of elution buffer (1%SDS, 100mM NaHCO_3_) at RT for 15 min. and collected. To reverse DNA-protein crosslinks NaCl was added to final concentration of 200mM and incubated overnight at 65ºC. RNA contamination was eliminated by treatment with RNAse A for 30 min. at 37ºC and proteins were cleared by Proteinase K at 45ºC for 2 hours. DNA was purified using QIAquick PCR Purification Kit (Quiagen, ID 28106) and eluted in 50 μl of DNAse-free Milli-Q water. Inputs, control IgG and anti-zic2 ChIP were sent to Fasteris SA (Switzerland) for quality test, library preparation and sequence service.
