## Supplementary material for "Zic2 abrogates an alternative Wnt signaling pathway to convert axon attraction into repulsion": Figure Legends to Supplemental Figures

**SUPPLEMENTAL FIGURE LEGENDS**

**Figure S1 related to Figure 1. Expression of different Wnts at the optic chiasm**

In situ hybridization in coronal sections of E16.5 embryos for different Wnts expressed at the optic chiasms.

**Figure S2 related to Figure 4. Transcriptomic profiles of known Zic2-target genes in Zic2 RGCs.**

**(A)** Pearson correlation matrix between normalized samples clustered by Euclidian dendrogram. EGFP-electroporated retinas: n=3; Zic2-electroporated retinas: n=3, biologically independent samples. **(B-D)** RNA-seq profiles for Zic2 (B), three well-known markers of iRGCs (C) and two markers of cRGCs **(D).**

**Figure S3 related to Figure 5. Zic2 ChIP-seq samples analysis**

**(A)** Left: Scheme summarizing the results of the Zic2 Chip-seq screen for RGC-specific (blue) and dSCNs-specific (red) targets. Right: Heat maps showing Zic2 binding at TSSs in the chromatin of RGCs (Retina) and dSCNs (SC). Intensity ranges from strong (blue) to weak (white). The signal is shown +/- 5kb from the TSSs. **(B)** Zic2 binding profile in the *EphB1* and *EphA4* loci in RGCs and dSCNs. **(C)** GO Biological process and Panther pathways enrichment analysis for genes associated with Zic2 peaks. **(D)** RNA-seq and ChIP-seq profiles for Zic2-bound genes related to the Wnt pathway that do not show significant changes in transcripts levels. **(E)** Percentage of upregulated and downregulated genes among Zic2 direct targets (common gene set). **(F)** Distribution of Zic2 binding across gene features for the two groups defined in panel E. **(G)** Distribution of motifs A and B the two groups defined in panel E.
